# Chromatin condensates tune nuclear mechano-sensing in Kabuki Syndrome by constraining cGAS activation

**DOI:** 10.1101/2024.05.06.592652

**Authors:** Sarah D’Annunzio, Lucia Santomaso, Daniela Michelatti, Chiara Bernardis, Giulia Vitali, Sara Lago, Claudia Testi, Emanuele Pontecorvo, Alessandro Poli, Fabrizio Pennacchio, Paolo Maiuri, Elodie Sanchez, David Genevieve, Lorenzo Petrolli, Thomas Tarenzi, Roberto Menichetti, Raffaello Potestio, Giancarlo Ruocco, Alessio Zippo

## Abstract

Cells and tissue integrity is constantly challenged by the necessity to adapt and respond to mechanical loads. Among the cellular components, the nucleus possesses mechano-sensing and mechanotransduction capabilities, yet the molecular mechanisms involved remain poorly defined. We postulated that the mechanical properties of the chromatin and its compartmentalization into condensates contribute to the nuclear adaptation to external forces, while preserving its integrity. By interrogating the effects of MLL4 loss-of-function in Kabuki Syndrome, we found that the balancing of transcriptional and Polycomb condensates tunes the nuclear responsiveness to external mechanical forces. We showed that MLL4 acts as a chromatin mechano-sensor by clustering into condensates through its Prion-like domain, and its response was regulated by the chromatin context. Furthermore, the mechano-sensing activity of MLL4 condensates is instrumental to withstand the physical challenges that nuclei experience during cell confinement and migration by preserving their integrity. In Kabuki Syndrome persistent rupture of nuclear envelope triggers cGAS-STING activation, which leads to programmed cell death. Ultimately, these results demonstrate the critical role chromatin compartments play in mechano-responses and how they impact pathological conditions by stimulating cGAS-STING signaling.

## Main

Tissue functionality and homeostasis rely on the capability of cells to respond and adapt to biochemical and mechanical environmental stimuli. The extracellular matrix (ECM) generates extrinsic mechanical forces which are transmitted and sensed by the cells through structural components and organelles, triggering different cellular responses over time and space^1, 2^. How biological systems encode and decode mechanical forces depends on the inherent physical properties of the cellular components, with the nucleus being constantly exposed to inward and outwards forces that can lead to its transient deformation while maintaining the genome integrity^3^. The ECM-mediated forces are directly transmitted through the cytoskeleton and LINC complex to the nuclear envelope (NE) and the nuclear lamina, further propagating the mechanical load to the chromatin. The nuclear lamina is a meshwork of the intermediate filament proteins Lamin A/C and B, whose relative abundance scales with the stiffness of the ECM^4, 5^. The nuclear mechanics is determined by the physical properties of the nuclear lamina and chromatin, which can modulate the magnitude and the time frame response to mechanical loads^6–10^. As chromatin behaves as a dynamic viscoelastic polymer in response to nuclear deformations, its organization is temporally reshaped, with an increment of euchromatin (H3 acetylation and H3K4 methylation) paralleled by a decrease of constitutive heterochromatin (H3K9me3), which is necessary to dissipate mechanical energy to ensure the genome integrity^11–14^. Albeit this evidence, the possible contribution of chromatin organization in determining the nuclear mechanical properties still needs to be understood.

Balancing inward and outward mechanical forces safeguards the maintenance of nuclear integrity. Indeed, it has been shown that during cell migration through confined environments, nuclear deformation leads to transient and localized NE rupture^15–17^. By nucleating the chromatin binding protein BAF, which is tethered by transient cytoplasmic exposure to genomic DNA, dedicated repair machinery comprising endosomal sorting complexes required for transport (ESCRT)-III, Lamins and LEM-associated proteins are recruited to the sites of NE rupture^17–21^. The persistence of NE rupture, which is frequently associated with pathological conditions compromising either the functionality or the protein abundance of Lamin A/C^17, 22–26^, triggers DNA damage and the cGAS-STING pathway, an innate immune signaling activated by cytosolic dsDNA^16, 17, 27, 28^. Indeed, it has been reported that in many physiological and pathological conditions, the recurrent loss of NE integrity initiates the cGAS-STING cascade, which leads to a diversity of cellular functions, including type I interferon response, inflammasome activation, and cell death pathways^29–31^. The contribution of the nuclear fragility in determining which kind of cellular response will be activated downstream to cGAS-STING pathway has yet to be determined. Nevertheless, many pathological conditions, including cancer, fibrosis, cardiomyopathies, and developmental disorders are related to alterations of the nuclear components involved in the mechano-transmission and mechano-responses^32–34^. Kabuki Syndrome (KS) is a rare genetic disease caused by the haploinsufficiency of the chromatin modifiers MLL4 (encoded by *KMT2D*) or UTX (encoded by *KMD6A*), whose clinical manifestations recall an altered mechano-signaling during cell lineage commitment and tissue homeostasis^35–37^. Both these chromatin regulators harbor intrinsic disorder regions (IDRs), which contribute to their compartmentalization into transcriptional condensates and counterbalance the repressive function of the Polycomb (PcG) complexes^32 38^. Besides determining the local chromatin environment to regulate gene expression, condensates are also involved in defining the 3D genome organization and the chromatin’s mechanical properties. In this respect, we have found that in KS the haploinsufficiency of MLL4 alters the equilibrium between transcriptional and PcG condensates, affecting the nuclear architecture and mechanics and cell lineage commitment^10 33^. Albeit their relevance, the mechanisms by which chromatin senses the mechanical forces and decodes this information are far from being understood.

We hypothesize that chromatin condensates may contribute to nuclear mechano-sensing and mechano-response. The assembly of condensates is governed by heterotypic interactions, which compartmentalize chromatin into functional domains. In this context, externally applied forces and the resulting nuclear deformation may be sensed by the chromatin condensates and decoded through their transient reorganization, thereby regulating the genome topology. To verify this hypothesis, we measured the response of transcription- and PcG-associated chromatin condensates to external mechanical cues in the context of KS. We found that MLL4 mediates the mechanical response by driving the nucleation of transcription condensates at the expense of PcG clustering. This imbalance results in chromatin softening, which secures nuclear integrity and prevents cGAS overactivation.

## Results

### Mechanical stimuli unbalance the transcription- and PcG-associated chromatin condensates

To address the contribution of chromatin condensates to external mechanical stimuli, we monitored the responses of mesenchymal stem cells (MSCs) to the increase of the ECM substrate stiffness in the genetic context of KS. Indeed, we adopted the disease model system in which the insertion of a *KMT2D* truncating mutation (MLL4^Q4092X^) causes MLL4 loss-of-function (LoF), impairing nuclear architecture and chromatin condensates balancing^10^. Single-cell analyses by quantitative immunofluorescence (Extended Data Fig. 1a) showed that *KMT2D* haploinsufficiency caused a reduction of MLL4 protein abundance irrespective of the substrate stiffness yet maintaining the puncta-like nuclear distribution. (Extended Data Fig. 1b). By analyzing the clustering of the transcription-associated cofactors we found that the increase of substrate stiffness was mirrored by an augmented intensity of BRD4 condensates, yet this association was not maintained upon MLL4 LoF (Fig. 1a). Of importance, this pattern was recapitulated employing an independent MSC clone carrying a different truncating mutation of MLL4 (encoding MLL4^P4093X^), and in KS patient-derived fibroblasts (Fig. 1a and Extended Data Fig. 1c-e). To rule out whether the ECM-transmitted mechanical forces control the organization of transcription condensates in dependency of MLL4 abundance, we performed super-resolution imaging by SIM microscopy on the MSCs maintained on a soft and stiff substrate, respectively. Super-resolution imaging by SIM revealed that BRD4 is distributed throughout the nucleus in clusters ranging from ∼0.03 μm^2^ to ∼0.75 μm^2^. Notably, MLL4 LoF reduced BRD4 clustering, resulting in fewer and smaller condensates in response to substrate stiffness (Fig. 1b). These results suggest that the increment of the external mechanical forces is associated with an augmented MLL4-dependent assembly of transcriptional condensates.

**Fig. 1.**
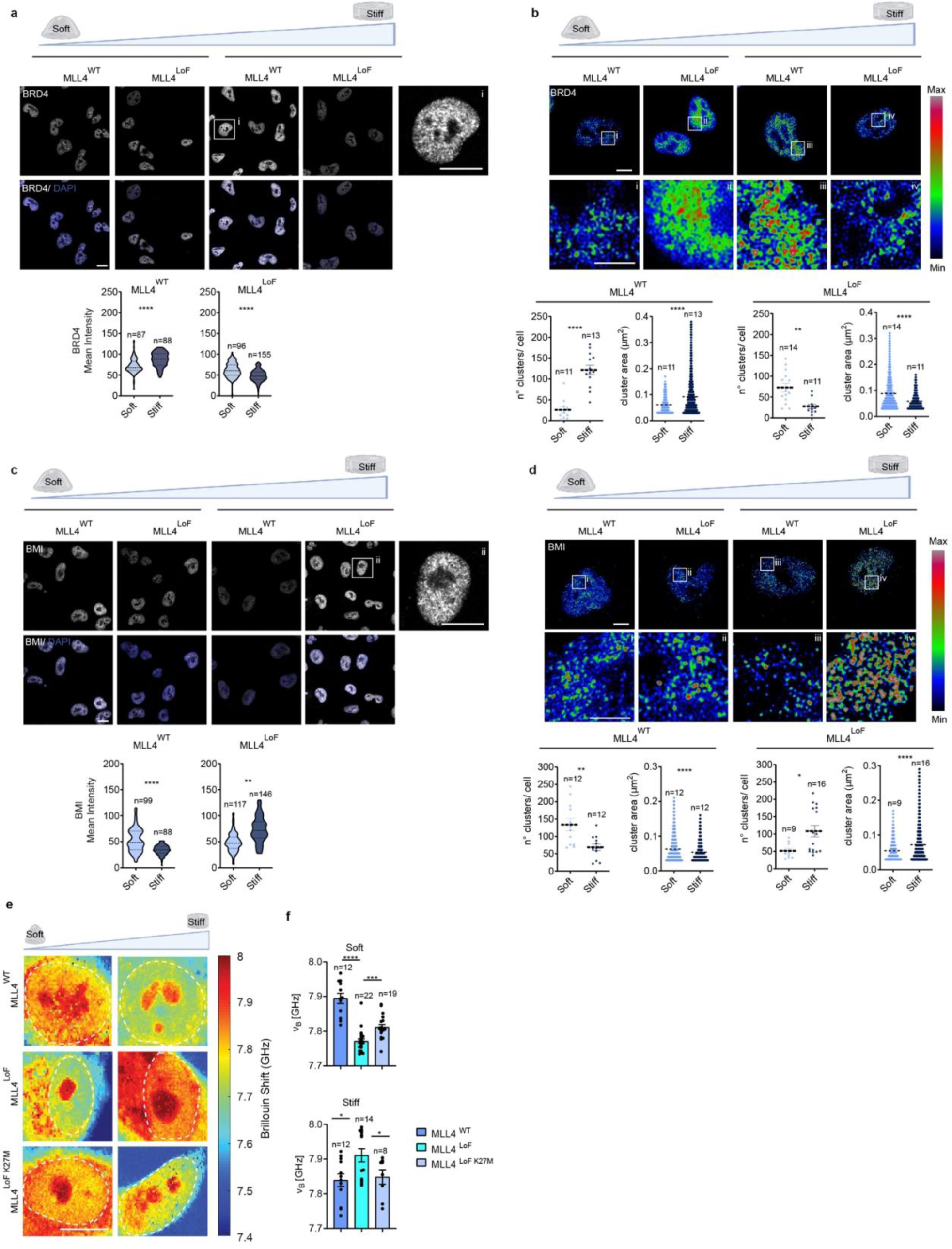
**(a)** Representative confocal images and relative quantifications of nuclear BRD4 mean intensity in MLL4^WT^ and MLL4^LoF^ MSCs on soft (0.5kPA) and stiff (32kPA) matrix. Scale bars, 10 µm. **(b)** Representative SIM images and relative cluster quantifications of immunostaining for BRD4 in MLL4^WT^ and MLL4^LoF^ MSCs, on soft and stiff matrix, scale bars 10 µm. Crops of BRD4 clusters are highlighted (i-iv), scale bars 5 µm. **(c)** Representative confocal images and relative quantifications of immunostaining for BMI in MLL4^WT^ and MLL4^LoF^ MSCs on soft and stiff matrix, scale bars 10 µm. **(d)** Representative SIM images and relative cluster quantifications of immunostaining for BMI in MLL4^WT^ and MLL4^LoF^ MSCs, on soft and stiff matrix, scale bars 10 µm. Crops of BMI clusters are highlighted (i-iv), scale bars 5 µm. **(e)** Representative maps of the stiffness distribution in MLL4^WT^ and MLL4^LoF^ MSCs, as well as MLL4^LoF^ MSCs expressing H3.3K27M, acquired with the Brillouin microscope. A higher Brillouin shift in the GHz frequency corresponds to a stiffer region of the cell. The dashed white lines enclose nuclei. Scale bars, 10 µm **(f)** Bar plots representing the nuclear Brillouin shift in WT and MLL4^LoF^. Scatter dot plots in **(b)** and **(d)** indicate Mean + S.E.M. Violin plots in **(a)** and **(c)** indicate median values (middle lines), and first and third quartiles (dashed lines). Bar plots in **(f)** show Mean + SEM. The number of cells (“n”) analyzed is reported in each panel. Statistical significance was determined by a two-tailed unpaired student’s t-test.

Considering that MLL4 LoF affected the clustering of PcG complexes^10^, we determined whether these repressive compartments were similarly responsive to the mechanical cues. Quantitative immunofluorescence analyses showed that although the transcription levels were unaltered (Extended Data Fig. 1f), clustering of the PRC1 components BMI1 and RING1B was diminished with the increment of the substrate stiffness (Fig. 1c and Extended Data Fig. 1g). Of note, the responsiveness of PcG clusters to the external mechanical forces was impaired in MSCs carrying *KMT2D* truncating mutations, and in KS patient-derived fibroblasts (Fig. 1c and Extended Data Fig. 1h-j). SIM microscopy indicated that the differences in PcG spatial organization were associated with a decrease in the abundance and the dimension of the condensates (Fig. 1d).

To rule out the contribution of PcG clustering to the altered responsiveness of the MLL4 ^Q4092X^ MSCs, we rescued its functionality by overexpressing the dominant-negative H3.3 K27M (Extended Data Fig. 1k)^10^. We found that by restoring PcG clustering, we re-established the relative abundance of the transcriptional condensates, both on soft and stiff conditions (Extended Data Fig. 1i). These results suggest that the balancing between transcription and PcG condensates could be involved in cellular mechano-response by mediating the nuclear architecture and mechanical properties. Indeed, by quantifying the longitudinal elastic modulus calculated on the base of the Brillouin frequency shift^39^ we found that the chromatin stiffness mirrored the abundance of PcG condensates (Fig 1e, f). This pattern was reversed upon MLL4 LoF and was rescued by re-establishing PcG condensates by expressing the H3.3K27M mutant. In sum, the depicted organization of chromatin condensates suggests that *KMT2D* haploinsufficiency impairs the nuclear mechano-response to ECM-mediated forces, strengthening the role of MLL4 in tuning the nuclear mechanical properties. These results indicate that chromatin condensates are responsive to external mechanical forces and their balancing possibly tune the nuclear mechanical properties.

### MLL4 abundance determines nuclear mechanical response to tissue stiffening

Next, we asked whether the measured increment of MLL4 clustering in response to mechanical forces also occurs in vivo and what its role is in tissue mechano-response. We first analyzed the abundance and the nuclear distribution of MLL4 along mice’s growth plate located at the epiphysis of long bones, which is the tissue site in which bone elongation takes place by endochondral ossification^40^. Of importance, immunofluorescence analyses showed that MLL4 was distributed in clusters within the nuclei of differentiating hypertrophic chondrocytes and osteoblasts while showing a low level of expression in the resting and proliferating chondrocytes that reside within the apical zone (Fig. 2a). Thereafter, we assessed the contribution of MLL4 to the nuclear mechanics of chondrocytes and osteoblasts, whose function and homeostasis are influenced by ECM-mediated mechanical forces^41^. To this end, we generated a KS mouse model (thereafter named *Kmt2d^cHe^*^t^) by crossing homozygous *Kmt2d* conditional allele mice *(Kmt2d^fl/fl^)* with CMV-Cre transgene^42^. By monitoring the body length and weight, we found that *Kmt2d^cHe^*^t^ mice showed a marked postnatal growth delay with respect to the *Kmt2d^fl/+^* counterpart (Fig. 2b and Extended Data Fig. 2a, b). Given this phenotype that resamples the postnatal growth delay affecting KS individuals^37^, we focused on measuring possible alterations in the morphology and tissue mechanics of the epiphysis of long bones. We noticed an enlargement of the growth plate and a poorly defined hypertrophic zone in the *Kmt2d^cHe^*^t^ condition compared to the *Kmt2d^fl/+^* littermates (Fig. 2c). Brillouin microscopy showed that the ECM stiffness increased longitudinally along the growth plate, showing the highest rigidity at the level of the primary spongiosa, the site for cartilage mineralization of the newly elongating bones (Fig. 2d, e). In the same condition, we found that the chromatin stiffening anti-correlated with the ECM rigidity, showing a lower level of stiffness in the osteoblasts residing in the spongiosa, with respect to the chondrocytes, in line with what we observed in vitro (Fig. 1g). Of importance, osteoblasts of the *Kmt2d^cHe^*^t^ mice showed a relative increment of nuclear stiffness, while the ECM mechanical properties were unperturbed. On the contrary, within the hypertrophic zone of the growth plate of the *Kmt2d^cHe^*^t^ mice, which is characterized by a modest ECM stiffness, we measured an augmented nuclear stiffening (Fig. 2d, e). Remarkably, the same change in nuclear mechanical properties was not detected within the proliferative zone, which is characterized by a low level of expression of MLL4 as well as by an increment of ECM stiffening in the *Kmt2d^cHe^*^t^ mice (Fig. 2d, e). In sum, the depicted organization of chromatin condensates suggests that *KMT2D* haploinsufficiency impairs the nuclear mechano-response to ECM-mediated forces, strengthening the role of MLL4 in tuning the nuclear mechanical properties.

**Fig. 2.**
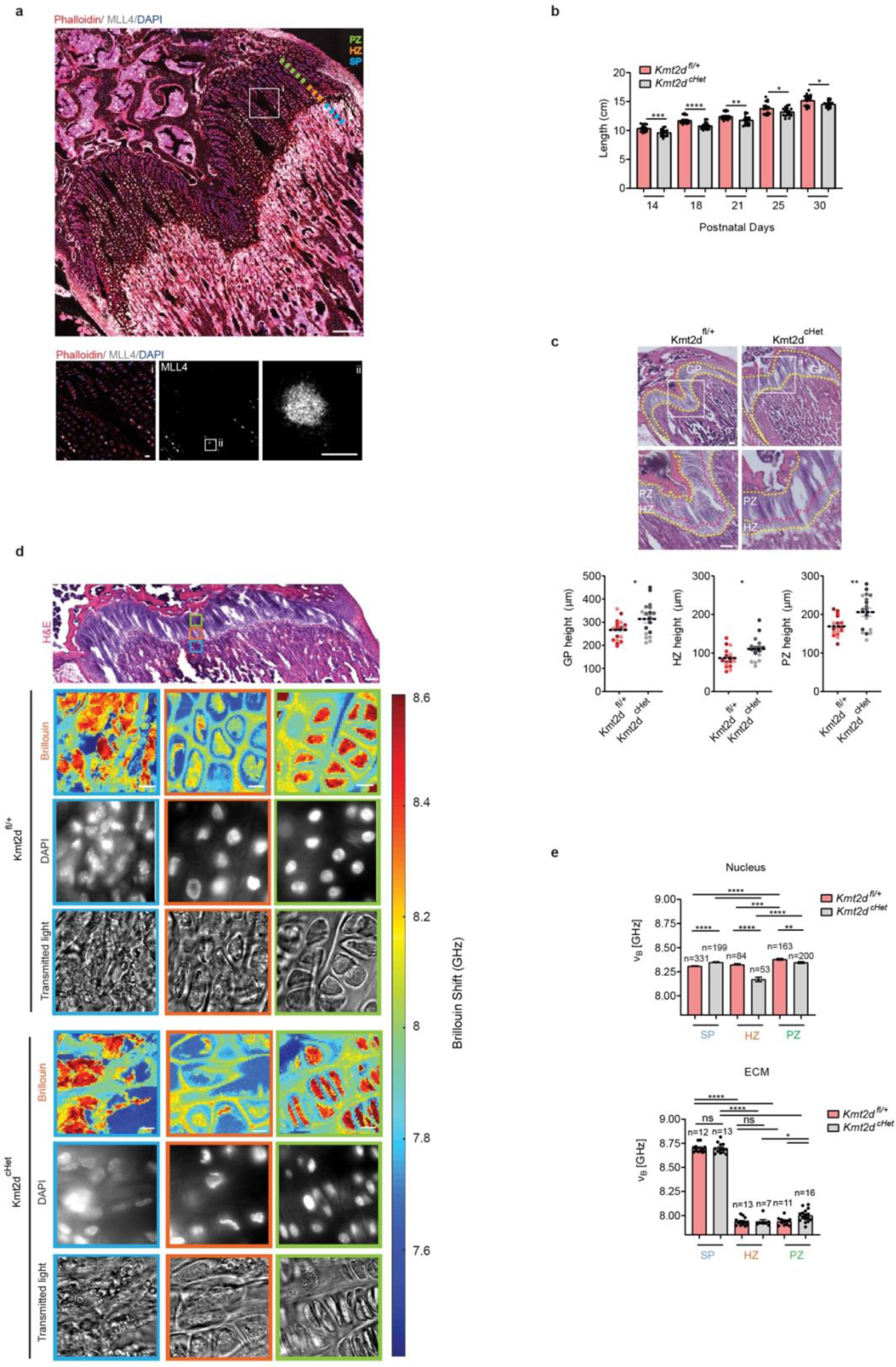
**(a)** Quantification of the length (cm) of Kmt2d^fl/+^ (WT) and Kmt2d^cHet^ (KS) at the indicated postnatal days (*n* = 18 mice derived from three different litters). **(b)** Representative images of MLL4 (white)/Phalloidin (red) immunostaining, scale bars 100 µm. Lower panels: (i) enlargement of MLL4/Phalloidin immunostaining in hypertrophic and proliferative cells within the growth plate, (ii) single-cell crop showing MLL4 clusters. Scale bars 10 µm. **(c)** Hematoxylin and Eosin (H&E) staining of Kmt2d^fl/+^ and Kmt2d^cHet^ femoral growth plate (GP), highlighted by a yellow dotted line. Within the GP the hypertrophic and proliferative zone (HZ and PZ, respectively) are marked. Below, quantification of GP, HZ and PZ height (*n* = 18, data combined from three mice in each group). **(d)** Upper panel: representative image of H&E staining of the femur growth plate in Kmt2d^fl/+^ (WT). Within the GP are highlighted the SP in blue, the HZ in green and the PZ in orange. Scale bars, 100µm. Below, representative maps of the stiffness distribution in the aforementioned regions, in Kmt2d^fl/+^ (WT) and Kmt2d^cHet^ (KS) GP. Scale bars, 10µm. The DAPI and transmitted light signal are also included. **(e)** Quantification of nuclear (upper panel) and ECM (lower panel) stiffness (Brillouin shift) in primary spongiosa (SP), hypertrophic zone (HZ) and proliferative zone (PZ) in Kmt2d^fl/+^ (WT) and Kmt2d^cHet^ (KS). The number of cells (upper panel) and ROI (lower panel) analyzed for each zone is reported as “n”. Scatter dot plots in **(c)** and bar plots in **(a)** and**(e)** indicate Mean + S.E.M. Statistical significance was determined by a two-tailed unpaired student’s t-test.

### MLL4 condensates are responsive to external mechanical forces

To define the possible mechanism by which transcriptional condensates sense the forces transmitted from the cytoskeleton to the nucleus, we measured the assembly of MLL4 condensates in living cells by adopting the light-activated optoIDR approach^43, 10^ (Fig. 3a). A single pulse of blue light-induced clustering of MLL4_PrLD giving rise into spherical droplets whose numerosity was independent from the protein abundance, while their dimension decreased linearly, as expected (Extended Data Fig. 3a, b). Time-lapse imaging showed that the optoMLL4 nucleated in droplets that, with time, coalesced into larger condensates (Fig. 3b-d). Of importance, this response resulted in being sensitive to the external mechanical forces, as we measured a reduced assembly of MLL4-PrLD clusters in MSCs grown on the soft matrix. In the same setting, we also determined an increment of the cluster size with respect to what we measured by keeping cells on the stiff matrix (Fig. 3b-d). We thus evaluated the contribution of the chromatin context to the dynamic clustering of optoMLL4, in both these mechanical regimes. We found that the assembly of the optoMLL4 in response to external mechanical forces was impaired in MLL4 ^Q4092X^ MSCs, giving rise to a reduced number of smaller condensates (Fig. 3b-d). Notably, by restoring the abundance of PcG condensates by means of expressing the H3.3 K27M mutant, we re-established the responsiveness of the optoMLL4 clusters to the mechanical load (Extended Data Fig. 3c). Altogether, these findings suggest that substrate rigidity affects the kinetics of MLL4-PrLD clustering in living cells. Furthermore, MLL4 LoF partially impairs condensate formation in response to external mechanical forces.

**Fig. 3.**
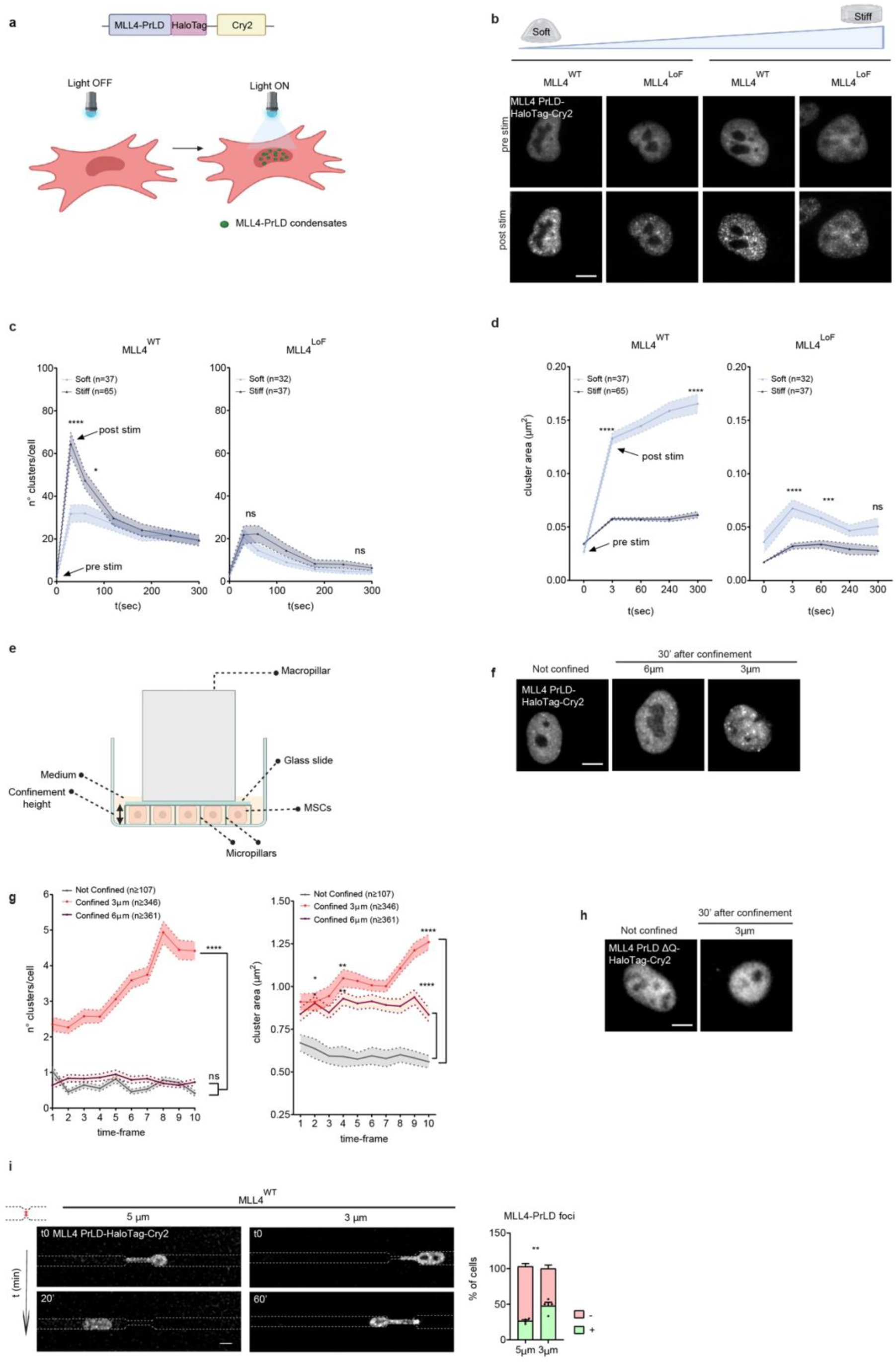
**(a)** Schematic representation of the MLL4-PrLD-HaloTag-Cry2 optogenetic tool. **(b)** Representative images of blue-light-induced clustering of MLL4-PrLD-HaloTag-Cry2 before and after the stimulus in MLL4^WT^ and MLL4^LoF^ MSCs on soft and stiff matrix. The cells were stimulated for 3 s with 488-nm light, followed by image acquisition. Scale bars, 5 µm. **(c-d)** Quantification of the number **(c)** and area **(d)** of light-induced droplets of MLL4-PrLD-HaloTag-Cry2 at different time points. The time point t=0 represents the pre-stimulus. **(e)** Schematic representation of the Cell Confinement. The height of confinement is controlled by polydimethylsiloxane (PDMS) micropillars, fabricated in a glass slide and attached to a PDMS piston. **(f)** Representative images of MLL4^WT^ MSCs over-expressing MLL4-PrLD-HaloTag-Cry2 confined at 3 and 6μm and the relative control (not confined cells). Scale bars, 5 µm. **(g)** Quantification of the number and area of MLL4-PrLD-HaloTag-Cry2 clusters at different time points in MLL4^WT^ MSCs not confined and confined at 3 and 6μm. **(h)** Representative images of MLL4-PrLD-ΔQ-HaloTag-Cry2 in NIH3T3 cells confined at 3μm and the relative control (not confined cells). Scale bars, 5 µm. **(i)** Representative images of MLL4^WT^ MSCs over-expressing MLL4-PrLD-HaloTag-Cry2 passing through constrictions of 5 and 3μm at different time points. On the right, quantification of the percentage of cells that spontaneously form (+) or do not form (−) MLL4-PrLD foci (*n* = 4 biologically independent samples). Scale bars, 5 µm. Scatter XY plots in **(c)**, **(d)**, **(g)** and Bar plots in **(i)** show mean + S.E.M. Statistical significance was determined by two-way ANOVA test for panels **(c)**, **(d)**, **(g)**, and by a two-tailed unpaired student’s t-test for panel **(i)**.

To determine whether the PrLD of MLL4 acts as a chromatin mechano-sensor, we measured the dynamic assembly of MLL4 condensates in response to applied mechanical load through nuclear confinement (Fig. 3e), independently from light-induced (CRY2 dependent) nucleation (Extended Data Fig. 3d). We observed that while both wild-type and MLL4^Q4092X^ MSCs responded to the increased confinement by activating the actin fibers, the MLL4 LoF caused a substantial increase in cell death at the most stringent confined condition (Extended Data Fig. 3e). Moreover, we found that by increasing the cell confinement MLL4-PrLD clustered in spherical droplets whose abundance and size increased linearly with time (Fig. 3f, g, Extended Data Fig. 3f, and Supplementary Video 1). Of importance, a very similar mechano-response was depicted by monitoring the clustering of the endogenous MLL4 in MSCs (Extended Data Fig. 3g). Given the central role of the polyQ tract in MLL4 clustering^10^, we determined its contribution to the mechano-response. We found that removing the polyQ stretch impaired the assembly of MLL4-PrLD condensates upon cell confinement (Fig. 3h). We finally assessed the clustering of MLL4-PrLD upon migration of MSCs throughout microchannels with restrictions, which imposes a transient mechanical load on the nucleus. Accordingly, we observed a linear increase in MLL4-PrLD clustering proportionally to the augmented level of mechanical load (Fig. 3i and Supplementary Video 2a, b). These results indicate that MLL4 can dynamically assemble into chromatin condensates in response to nuclear mechanical load, possibly acting as a chromatin mechano-sensor.

### Mechanical forces guide the folding transition of the polyQ stretch of MLL4-PrLD

As the mechano-signaling relies on the capability of macromolecules to sense and transmit the received signals by transiently changing their conformations^44, 45^, we investigated whether the PrLD of MLL4 can go through a folding transition in response to mechanical load. The prediction of the MLL4-PrLD structure highlighted that despite belonging to an intrinsically disordered domain of the MLL4 sequence, the embedded polyQ tract is associated with an inherent likelihood to yield locally folded structures such as α-helixes (Fig. 4a, b). This observation corroborated the choice of this region as the starting coordinates of the system employed in the molecular dynamics setup. Indeed, to establish the role of the polyQ tract in the clustering of MLL4 from a mechanistic perspective, we carried out molecular dynamics (MD) simulations enforcing increasingly higher target pressures on a single polyQ moiety.

**Fig. 4.**
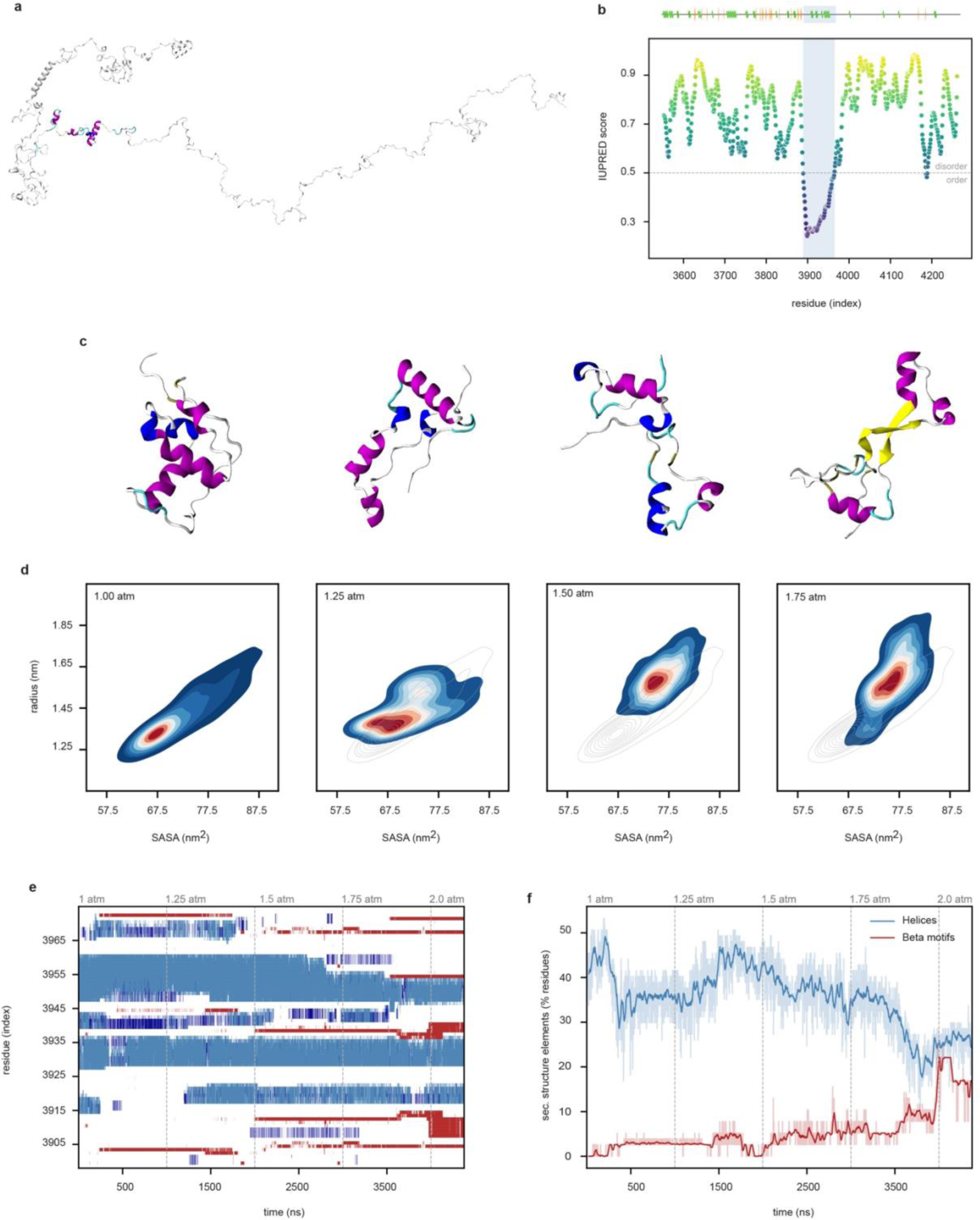
**(a)** Structure prediction of the Prion-like domain of MLL4 (MLL4-PrLD, residues 3561 to 4270 of the MLL4 sequence); the polyQ tract at the core of MLL4-PrLD (residues 3898 to 3974) is highlighted. **(b)** Top: Layout of the secondary structure motifs associated with the structural prediction of MLL4-PrLD by the IntFold web server: Helices and beta strands are shown as coils and arrows, respectively, and the polyQ tract of MLL4-PrLD (residues 3898 to 3974) is highlighted (shaded area). Illustration obtained *via* the Biotite Python toolkit. Bottom: Prediction of structural disorder of the MLL4-PrLD based on the IUPRED bioinformatics tool: Lower IUPRED scores are associated with a higher tendency for each residue to fold into ordered structures. Likewise, the polyQ tract of MLL4-PrLD is highlighted (shaded area). **(c)** Snapshots of the structure of the polyQ tract of MLL4-PrLD from MD simulations, taken as the final frames of the 1-microsecond trajectories at 1 (leftmost), 1.25, 1.5, and 1.75 atm (rightmost), highlighting a consistent evolution in the secondary and tertiary structure, and the emergence of beta-like motifs - shown in yellow. **(d)** Putative free energy profiles of the polyQ tract of the Prion-like domain of MLL4 (MLL4-PrLD), along the twofold reaction coordinate defined by the radius of gyration and the solvent-accessible surface area (SASA). All subplots relate to a different target pressure of the molecular dynamics (MD) setup, whereby each value of P has been sampled for 1 microsecond. A color scheme defines the likelihood for the system to adopt specific values of the reaction coordinates throughout the MD trajectory, from blue (lowest) to red (highest frequency): A transition between two basins - towards an expanded conformation of the polyQ tract - is associated with the external pressure ramp. In fact, a decreasing trend of both reaction coordinates observed in the earliest frames at 1 atm (which is replicated within each subplot by dark gray contour lines) relates to an early equilibration stage of the system. **(e)** Left: Profile of the evolution of the secondary structure of the polyQ tract of MLL4-PrLD via MD simulations: Each residue is attributed a secondary structure motif as beta strands (red), alpha helices (cyan), alt. helices (blue), coils/turns (white) on a frame-wise basis along the trajectory - dashed vertical lines mark the stages of the pressure “ramp”. Right: Fraction of residues of the polyQ tract of MLL4-PrLD involved with either helices or beta motifs along the MD trajectory, on a frame-wise basis - likewise, dashed vertical lines mark the stages of the pressure ramp, whereas rolling averages of the signals are included as darker lines for clarity.

The observed dynamics of the sole polyQ tract feature a broad transition between conformational basins, whereby the layout of the system is driven from a “compact” to an “expanded” configuration (Fig. 4d and Extended Data Fig. 4a): This is mainly accounted for by a re-adjustment of the stable helical motifs in the sequence, i.e., conserved in over 75% of the trajectory frames and involving residues 3929-3936 and 3949-3955 of MLL4 (Fig. 4c).

Notably, this global conformational transition somewhat eases a steady drying-up of the helical content of the polyQ tract and the emergence of beta-like motifs (Fig. 4f) - the latter being sparsely distributed in the earliest stages of the MD trajectory, eventually yielding a stably-folded, extended beta sheet in an antiparallel arrangement (Fig. 4c, e). An increase in the external pressure seemingly enhances this process and involves residues that originally belong to both coiled and helical domains - that is, according to the starting configuration of the system (Fig. 4e). Moreover, the newly achieved ***β***-motif is conserved by further adjusting the target external pressure, as observed via a shorter 2 atm MD extension. Thus, molecular dynamics simulations suggest that the polyQ tract of MLL4-PrLD exhibits several conformational polymorphs and that increasing the external pressure seemingly stabilizes an extended, ***β***-rich configuration.

### Unbalancing chromatin condensates affects nuclear envelope tension in response to mechanical cues

IN light of these findings, we measured whether MLL4 LoF impaired the responsiveness of MSCs to the increment of mechanical forces. We first investigated the contribution of Lamin A/C to define the nuclear mechanics, considering that its abundance increases in response to ECM stiffness^5, 15^. We found that while in the wild-type cells Lamin A/C abundance increased in response to the augmented stiffening of the ECM, the *KMT2D* haploinsufficiency resulted in reduced responsiveness to the same mechanical load (Fig. 5a). Remarkably, the impaired responsiveness to ECM stiffness was further verified using an independent clone of MSCs, and in patient-derived fibroblasts (Extended Data Fig. 5a, b). Of note, a similar pattern was observed when we analyzed the level and distribution of H3K9me3, a marker of constitutive heterochromatin, which is frequently associated with the nuclear lamina^46^ (Extended Data Fig. 5c). By analyzing the nuclear morphology, we found that the nuclear area, volume, and flatness were increased in cells maintained on a stiff matrix, and this response was weakened upon MLL4 LoF (Fig. 5b, c and Extended Data Fig. 5d-e). Notably, in the same cellular condition, we re-established the nuclear shape and the relative abundance of Lamin A/C by rescuing the clustering of PcG condensates (Fig. 5d Extended Data Fig. 5f). By measuring the excess of nuclear perimeter, we found that the adjustment of the nuclear shape in response to the increment of mechanical forces was mirrored by an augmented NE tension (Fig. 5e and Extended Data Fig. 5g, h). The NE of *KMT2D* haplo-insufficient cells showed invaginations and a reduced nuclear perimeter, resembling the pattern observed in cells maintained on a soft matrix (Fig. 4e and Extended Data Fig. 5g, h). To rule out the contribution of the Lamin A/C decrease to the altered NE tension, we restored its abundance by expressing Lamin A-HaloTag in MLL4^Q4092X^ MSCs. In this setting, despite a comparable level of Lamin A/C, we did not observe any rescue of the NE surface (Extended Data Fig. 5i, j), indicating that the altered nuclear shape was not solely dependent on the relative abundance of A/C. To measure the NE tensional state of MSCs directly in response to ECM stiffness, we adopted the Mini Nesprin 1 reporter as the FRET sensor^47^. We found that while in the MLL4^WT^ MSCs the NE tension increased with the stiffening of the matrix, this responsiveness was impaired upon MLL4 LoF (Fig 5f). In sum, the obtained results indicate that the *KMT2D* haploinsufficiency reduces the NE responsiveness to the external mechanical load.

**Fig. 5.**
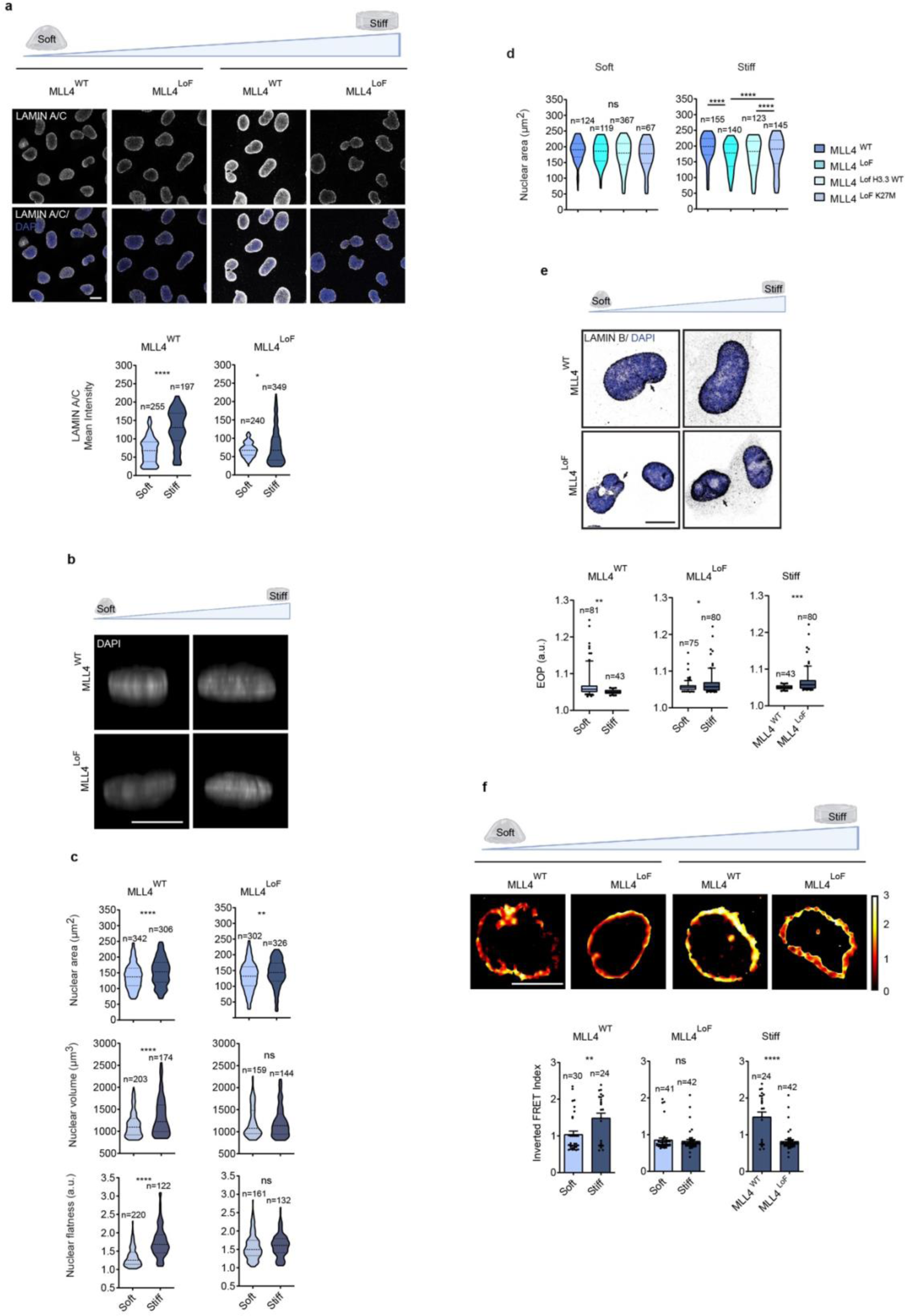
**(a)** Representative confocal images and relative quantifications of immunostaining for LAMIN A/C in MLL4^WT^ and MLL4^LoF^ MSCs on soft and stiff matrix. Scale bars, 10 µm. **(b)** 3D side view of MLL4^WT^ and MLL4^LoF^ MSCs nuclei stained with DAPI on soft and stiff matrix. Scale bars, 10 µm. **(c)** Quantification of nuclear volume, area, and flatness in MLL4^WT^ and MLL4^LoF^ MSCs on soft and stiff matrix. **(d)** Quantifications of the nuclear area in MLL4^WT^ and MLL4^LoF^ MSCs and MLL4^LoF^ MSCs expressing either H3.3WT or H3.3K27M on soft and stiff matrix **(e)** Representative confocal images of immunostaining for LAMIN B in MLL4^WT^ and MLL4^LoF^ MSCs on soft and stiff matrix. Black arrows indicate the presence of nuclear envelope invaginations, quantified as EOP (Excess Of Perimeter). **(f)** Representative images showing the FRET Efficiency of the Mini-Nesprin FRET sensor in MLL4^WT^ and MLL4^LoF^ MSCs on soft and stiff matrix. Below, a quantification of the Inverted FRET; lower values indicate decreased nuclear envelope tension. Violin plots in **(a)**, **(c)** and **(d)** indicate median values (middle lines), and first and third quartiles (dashed lines). Box plots in **(e)** indicate the median (middle line), the first and third quartiles (box), and the 10th and 90th percentile (error bars) of the EOP. Bar plots in **(f)** show Mean + SEM. The number of cells (“n”) analyzed is reported in each panel. Statistical significance was determined by a two-tailed unpaired student’s t-test.

### MLL4 LoF impairs nuclear envelope integrity upon mechanical stress

Considering that the MLL4-associated phenotype resembled the perturbation of the NE integrity that characterizes pathological conditions such as cancer and laminopathies^16, 17, 48^, we measured the frequency of NE rupture in response to ECM stiffening. To assess the frequency of spontaneous NE rupture, we quantify the clustering of the endogenous cGAS, which senses the exposure of chromatin to the cytoplasm^16, 17^ (Fig 6a). We found that ECM-mediated NE softening was mirrored by increased cGAS clustering in proximity to the NE. Of note, this pattern was resembled by the MLL4 LoF as both MSCs and KS patient-derived fibroblasts grown on a stiff matrix showed an increment of spontaneous NE rupture (Fig. 6a and Extended Data Fig. 6a, b). The increased NE fragility of MLL4^Q4092X^ MSCs was further verified by monitoring the colocalization of cGAS and BAF, which is involved in the repair of the NE ruptures^17, 19, 21, 28^. The redistribution of the BAF-mCherry in the proximity of the nuclear envelope occurred at the same sites in which cGAS clustered, irrespective of the source of NE fragility (Fig. 6b, c). To better determine the transient loss of NE integrity during nuclear deformation, we combined the NLS-GFP reporter system with the migration of MSCs through the confined environment of microchannels (Fig 6d). By monitoring the relative accumulation of the GFP signals in the cytoplasm after the nuclear engagement with the microchannel restrictions (Fig 6e, f and Supplementary Video 3a-c), we found that in wild-type MSCs the frequency of NE rupture increased with the augmented degree of confinement, in line with previous reports^17^ (Fig. 6g). Of importance, this response was altered in the MLL4 LoF condition, as the NE rupture occurred in all the cells encountering the least stringent confinement (Fig. 6g, Extended data Fig. 6c and Supplementary Video 3d-f). Notably, this pattern was rescued upon re-establishing the PcG condensates by means of expressing the H3.3K27M mutant (Extended data Fig. 6d, e). To determine whether this transient NE rupture culminates in the exposure of the genomic DNA in the cytoplasm and eventually in the activation of the NE repair machinery, we monitored the clustering of both cGAS and BAF in living cells. We found that shortly after passing through the constrictions, the cGAS-GFP biosensor clustered in foci, whose frequency increased proportionally to the degree of confinement (Fig. 6h, i, and Supplementary Video 4a-c). The MLL4^Q4092X^ MSCs showed an increased occurrence of cGAS clustering irrespective of the compressive forces on the nucleus (Fig 6h, I and Supplementary Video 4d-f). Of note, the events of NE rupture were often associated with mechanically induced DNA damage, as visualized by the accumulation of 53BP1 after the passage through the constrictions (Extended data Fig. 6f, g). Nevertheless, the MLL4^Q4092X^ MSCs did not show a higher frequency of DNA damage, thus suggesting that the genome stability is similarly preserved (Extended data Fig. 6f, g). As the efficacy of NE repair is associated with the abundance and the extent of cytosolic BAF accumulating at the rupture site^17, 19, 21, 28^, we monitored in living cells the dimensions of the BAF clusters at the NE rupture sites that were visualized by using the cGAS-GFP biosensor. We found that while the cGAS foci had a comparable size, the dimension of the BAF clusters was reduced in MLL4^Q4092X^ MSCs (Fig 6j). Together, these data indicate that in response to nuclear mechanical stress, the NE integrity is impaired in *KMT2*D haplo-insufficient cells.

**Fig. 6.**
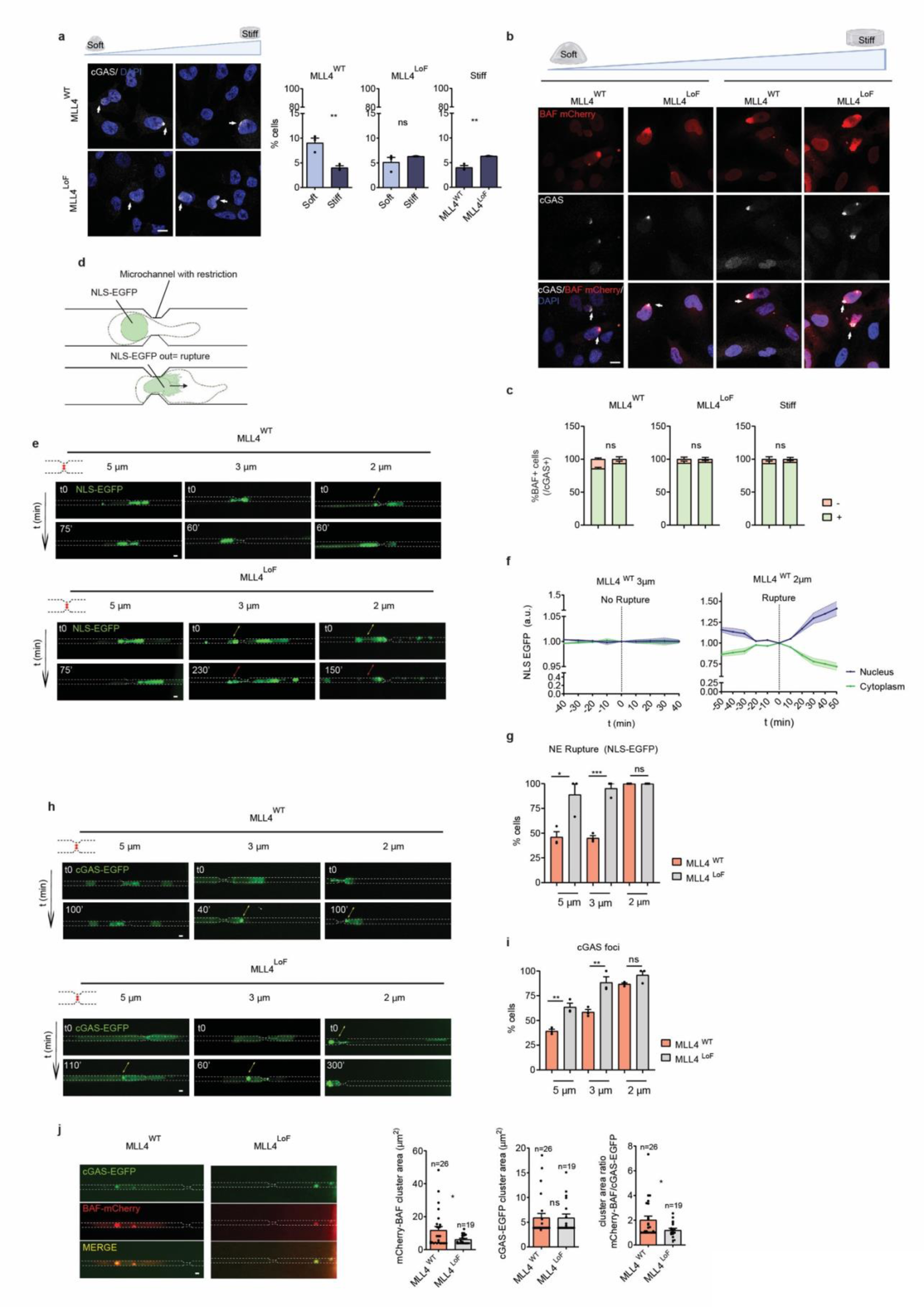
**(a)** Representative confocal images of immunostaining for cGAS in MLL4^WT^ and MLL4^LoF^ MSCs on soft and stiff matrix. Scale bars, 10 µm. On the right, quantification of the percentage of cells showing cGAS accumulation at the nuclear periphery, an indication of NE rupture (white arrows) (n=3 biologically independent samples). **(b)** Representative confocal images of immunostaining for cGAS (white) in MLL4^WT^ and MLL4^LoF^ MSCs over-expressing BAF-mCherry (red) on soft and stiff matrix. Scale bars, 10 µm. **(c)** Quantification of the percentage of cGAS positive cells showing also BAF accumulation at the nuclear periphery in correspondence of the cGAS foci (colocalization) (n=3 biologically independent samples). **(d)** Schematic representation of the NLS-EGFP reporter for measuring NE rupture. When the nucleus ruptures the NLS-EGFP signal (green, in the nucleus) diffuses into the cytoplasm (out= rupture). **(e)** Representative images of MLL4^WT^ and MLL4^LoF^ MSCs expressing NLS-EGFP passing through constrictions of different widths (5, 3, and 2 μm). Yellow arrows indicate cells that are subjected to NE rupture, and red arrows indicate cell death. Scale bars, 10 µm. **(f)** NLS-EGFP normalized mean intensity inside the nucleus (blue) and in the cytoplasm (green) of MLL4^WT^ in case of NE rupture (2 μm) or not (3 μm) (n≥4). Cytoplasmatic NLS-EGFP intensity was normalized to initial nuclear intensity and vice-versa to obtain NLS nuclear or cytosolic signal. Time equal to 0 (x axis) corresponds to the tip of the nucleus reaching the end of the constriction in which NE rupture occurs. **(g)** Bar plots showing the % of cells undergoing NE rupture in NLS EGFP-MSCs (n=3 biologically independent samples)**. (h)** Representative images of MLL4^WT^ and MLL4^LoF^ MSCs expressing cGAS-EGFP passing through constrictions of different widths (5, 3, and 2 μm). Scale bars, 10 µm. **(i)** Bar plots showing the % of cells showing cGAS foci in cGAS EGFP-MSCs (n=3 biologically independent samples). **(j)** Representative images of MLL4^WT^ and MLL4^LoF^ MSCs co-expressing cGAS-EGFP and BAF-mCherry passing through constrictions of 3μm. Scale bars, 10 µm. On the right, quantification of the area of the cGAS-EGFP and BAF-mCherry foci (n=3 biologically independent samples). The number of cells (“n”) analyzed for each sample is reported in the figure. Bar plots in **(a)**, **(c), (g), (i)** and **(j)**, and Scatter XY plots in (f) show mean + SEM. Statistical significance was determined by a two-tailed unpaired student’s t-test.

### Loss of NE integrity drives the cGAS/STING-mediated cell death in KS

Besides nucleating the NE repair machinery, BAF also plays a central role in controlling the activation of the cGAS-STING pathway by restricting cGAS binding to the genome after NE rupture, preventing the assembly of DNA-cGAS oligomers and its enzymatic activity^28^. Although the cellular distribution of BAF was not altered in MLL4^Q4092X^ MSCs, we found that its protein level was decreased (Fig. 7a and Extended Data Fig. 7a-d), possibly contributing to the activation of the cGAS-STING pathway upon mechanical stress. To verify this, we monitored the activation of the downstream effector IRF3 and found that in response to NE rupture, it was translocated into the nucleus (Fig. 7b). With the increase of ECM-mediated mechanical load, MLL4^Q4092X^ MSCs showed a higher frequency of cGAS-STING pathway activation (Fig. 7b and Extended Data Fig. 7e). Interestingly, recapitulating the decrease in BAF1 abundance reflected in MLL4^Q4092X^ MSCs did not affect the cGAS STING pathway response in MLL4WT MSCs, although further reducing it mimicked the MLL4 LoF phenotype. (Extended Data Fig. 7f, g).

**Fig. 7.**
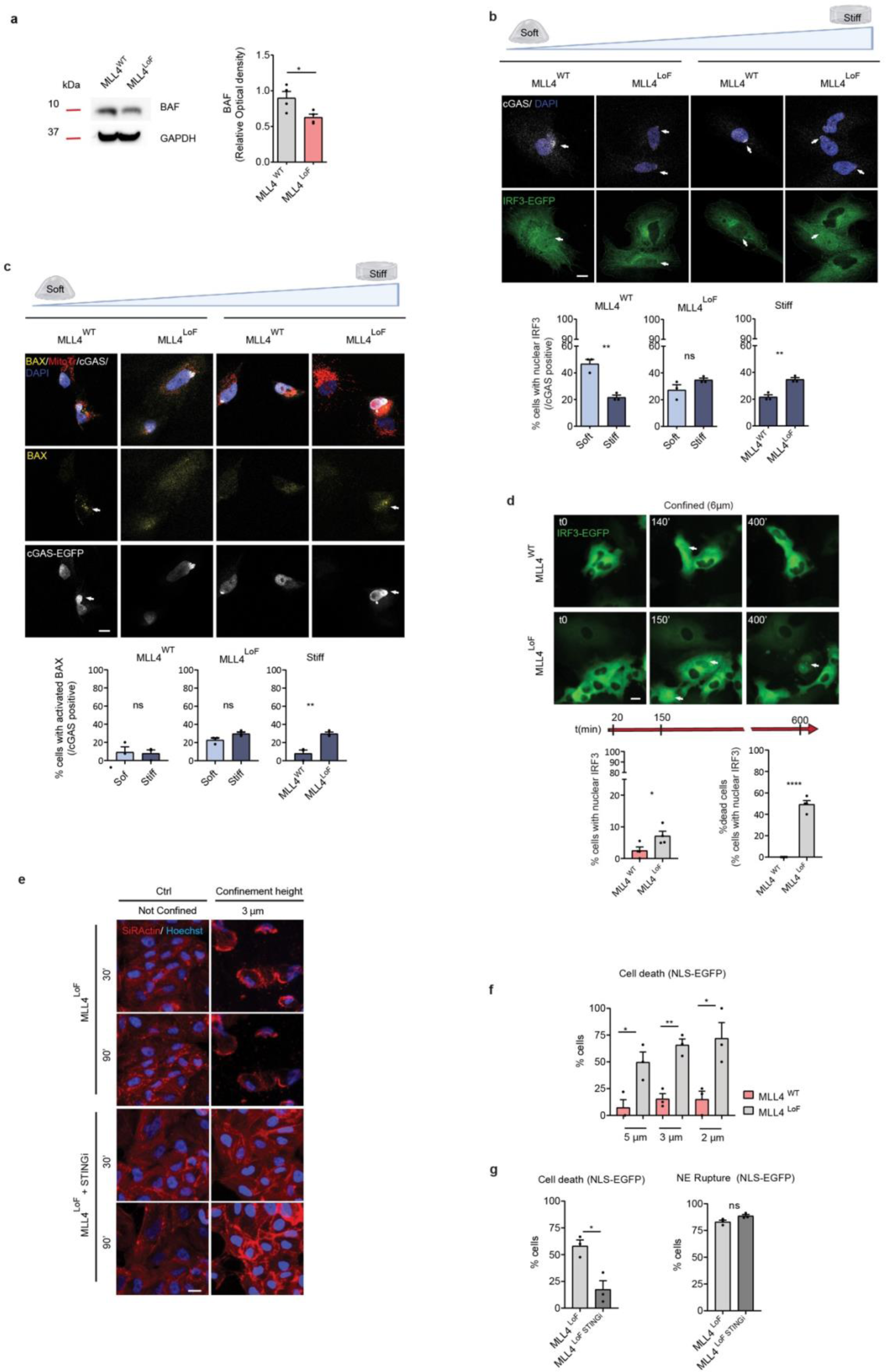
**(a)** Western Blot analysis of BAF in MLL4^WT^ and MLL4^LoF^ MSCs; GAPDH was used as a loading control. On the right: signal quantification for BAF normalized on GAPDH levels (n= 4 biologically independent samples). **(b)** Representative confocal images of immunostaining for cGAS in MLL4^WT^ and MLL4^LoF^ MSCs over-expressing IRF3-EGFP on soft and stiff matrix. Scale bars, 10 µm. Below is a quantification of the percentage of cells harboring IRF3 nuclear localization calculated over the cGAS positive cells (n=3 biologically independent samples). **(c)** Representative confocal images of immunostaining for BAX (yellow) in MLL4^WT^ and MLL4^LoF^ MSCs over-expressing cGAS-EGFP (white) on soft and stiff matrix. The Mito Tracker (red), was used to visualize mitochondria. Scale bars, 10 µm. Below is a quantification of the percentage of cells with activated BAX measured over the cGAS positive cells (n=3 biologically independent samples). **(d)** Representative images of MLL4^WT^ and MLL4^LoF^ MSCs over-expressing IRF3-EGFP under cell confinement of 6μm. Scale bars, 10 µm. Below is a quantification of the percentage of cells harboring IRF3 nuclear localization in the first 150’ of time-lapse. On those cells having nuclear IRF3, the percentage of cells that undergo cell death within 600’ of time-lapse was calculated (n=4 biologically independent samples). **(e)** Representative images showing MLL4^LoF^ MSCs treated or not with the STINGi not confined and under confinement of 3μm at the indicated time points. Cell nuclei are stained with Hoechst (blue), whether the cytoskeleton is marked in red (SirActin). Scale bars, 10 µm. **(f)** Bar plots showing the % of cells undergoing cell death in NLS EGFP-MSCs passing through restrictions of different widths (n=3 biologically independent samples)**. (g)** Bar plots showing the % of cells undergoing NE rupture and cell death in NLS EGFP-MSCs treated or not with the STINGi passing through restrictions of 3μm (n=3 three independent samples). Bar plots in **(a)**-**(d)** and **(f)-(g)** show mean + SEM. Statistical significance was determined by a two-tailed unpaired student’s t-test.

To figure out whether the cGAS-STING is primed by the increment of mechanical forces in a physiological context, we monitored the cGAS clustering along the growth plate and the primary spongiosa of long bones. We found that cGAS clustered in proximity to the NE of osteoblasts and osteoclasts residing in the primary spongiosa, which is characterized by the highest level of ECM stiffening (Extended Data Fig 7h and Fig. 2e. f). Of importance the frequency of cGAS activation resulted in being augmented in the *Kmt2d^cHe^*^t^ (Extended Data Fig 7h).

These data suggested that the cGAS-STING pathway is activated under mechanical stress in the KS pathological setting. Besides mediating IFN-dependent inflammatory response, cGAS-STING activates different evolutionarily ancient defense programs, including non-canonical autophagy and NLRP3-mediated inflammasome, which leads to different DNA-stimulated programmed cell death (PCD) pathways^49–52^. Indeed, we found that the executors of PCD pathways BAX/BAD^49, 53–55^ resulted activated in MLL4^Q4092X^ MSCs grown on stiff matrix (Fig. 7c and Extended Data Fig. 7i). By loading mechanical forces through cell confinement (Fig 3e), we measure a dynamic increase of cGAS-STING pathways, which culminated with 50% of cell death upon MLL4 LoF (Fig. 7d). To determine the contribution of chromatin clustering unbalancing in the activation of the pathway, we re-established the PcG condensates by expressing the H3.3K27M mutant in MLL4^Q4092X^ MSCs and found that the level of IRF3 activation was rescued, and the consequent PCD was strongly inhibited (Extended Data Fig. 7j). Using the most stringent confined conditions (3μm), we observed a rapid induction of PCD, which was abrogated by inhibiting STING (Fig. 7e). To verify that under mechanical stress the cGAS-STING axis activated PCD pathways upon MLL4 LoF, we monitored cell death of MSCs challenged by the passages through the microchannel restrictions. We found that while wild-type MSCs rarely underwent cell death after migrating through the confined environments, many *KMT2D* haploinsufficient cells dyed in response to mechanical stress (Fig. 7f and Extended Data Fig. 7k). This response was partially rescued by re-establishing the proper chromatin compartmentalization of PcG proteins (Extended Data Fig. 7l). At the same time, while treatment with STING inhibitor did not reduce the frequency of NE rupture, it interfered with PCD pathways, re-establishing the cell vitality of *KMT2D* haploinsufficient cells (Fig. 7g and Extended Data Fig. 7m). Together these findings indicated that in the KS condition, the increased frequency of NE rupture occurring in response to nuclear mechanical load, drove the activation of the cGAS-STING signaling, leading to PCD.

## Discussion

Genome compartmentalization into chromatin condensates has been viewed as a means of increasing the efficacy and specificity of biochemical processes, including transcription regulation and DNA repair. Here, we show that the balance between transcription and PcG condensates is instrumental for tuning the nuclear mechanics and preserving nuclear integrity in response to external stimuli. Indeed, we found that MLL4 LoF hampers nuclear responsiveness to external mechanical stimuli by reducing the assembly of transcriptional condensates. Of importance, the associated perturbation of the nuclear mechanical properties was restored by rescuing the clustering of PcG, indicating that the inward forces generated by chromatin compartmentalization are mechanically coupled with the outward mechanical stimuli. This observation, combined with recent findings underlying the contribution of chromatin folding in determining the nuclear mechanics, suggests that chromatin-associated proteins are capable of perceiving and responding to mechanical forces to withstand these physical challenges temporally. Here, we found that MLL4 can act as a chromatin mechano-sensor, by means of transiently clustering into condensates in response to mechanical load. Of note, this responsiveness is mediated by the MLL4-PrLD, as deleting the characterizing polyQ stretch significantly reduced the mechano-stimulated clustering. Considering that many mechano-sensors exert their activity by transiently changing their conformation^44, 45^ we simulated the folding of the PrLD, and we observed local transitions from α-helixes toward β-motifs in response to an increase int the external pressure applied. This response recalls what has been previously related to polyglutamine (polyQ) diseases in which the expanded polyQ stretch of the disease-causative proteins undergo a conformational transition from the monomer into the β-sheet-rich monomer that trigger the nucleation of soluble oligomers, which eventually age into amyloids^56, 57^. Further investigation would be required to demonstrate that the assembly of MLL4 condensates relies on the predicted transition to β-strand conformation. Nevertheless, we can not exclude that mechanical-related biochemical signaling associated with nuclear deformations may play a critical function in modulating chromatin compartmentalization^58–60^.

We also showed that the perturbation of condensates balancing predisposes to NE fragility that is exacerbated once cells are challenged by the necessity to reshape the nuclear morphology during cell migration and in response to confinement. These results underline the biological relevance of preserving nuclear integrity upon physical challenges, which otherwise may lead to persistent NE rupture. Indeed, we found that in the KS condition, the increment of nuclear deformation was mirrored by an augmented frequency of NE ruptures, leading to cGAS-STING activation that triggered PCD pathways. Sensing cytosolic DNA activates cGAS-STING, which stimulates a broad range of stress responses, including interferon signaling, non-canonical autophagy, or inflammasome activation that can eventually lead to cell death^49^. What factors guide the cellular choice of activating PCD pathways downstream to cGAS-STING signaling have yet to be determined. Possibly, the extent of the stimulus combined with the integration with other signaling pathways may determine the cellular outcome. In this respect, we observed that in the context of KS, the relative abundance of BAF1 contributes to the activation of cGAS-STING, indicating that multiple factors can tune the pathway and the consequent biological outcome. Further experiments would be required to define whether alterations of the chromatin organization can predispose cells to mount a more robust response upon exposure to chromatin in the cytoplasm because of NE rupture.

Of importance, many of the clinical manifestations of KS are attributable to alterations in the nuclear mechanics and integrity of the nuclear envelope. Indeed, many affected tissues rely on a proper response to mechanical stimuli that guide cell lineage commitment during development or control proper tissue homeostasis and functionality postnatally. The latter condition is properly exemplified by the postnatal growth delay affecting KS individuals, which we showed to correlate with an altered endochondral ossification with osteoblasts being characterized by a increase of chromatin stiffening that was mirrored by an augmented activation of cGAS-STING pathway. Whether a similar pattern contributes to determining other tissues’ functional alterations characterizing KS individuals warrants further investigation. Our findings underline the relevance of investigating the contribution of chromatin condensates in modulating nuclear mechanics and the downstream signaling, whose targeting may represent a therapeutic option for treating KS patients. Of interest, recent findings establish the cGAS–STING pathway as a driver of chronic neurodegenerative conditions. In this context, treatment with STINGi rescues physical and cognitive functions in several mouse models^61, 62^. Albeit in another pathological background, this evidence may further support the notion that inhibition of cGAS-STING could alleviate some of KS manifestations related to impairment of mechano-signaling.

## Acknowledgments

We thank the imaging facility from CIBIO for SIM microscopy and data analyses. We thank Prof. Fulvio Chiacchera and Elisa Ferracci for the fruitful scientific discussion and for histological sample preparation. We thank D. Allis for sharing the H3.3K27M construct, K. S. Zaret for TETO-FUW-LMNA-HALO vector, R. Young for the OptoIDR vectors, L. Tiberi forpCMV-HalyPBbase plasmid and M. S. Wiebe for providing BAF construct and antibody. We also thank I. Pastushenko for providing the Kmt2dfl/fl mouse strain. Maiuri group is supported by the Italian Association for Cancer Research (AIRC), IG#24976. Alessandro Poli’s work was founded by Fondazione Umberto Veronesi Post-doctoral fellowships (#000359) and Short-EMBO Fellowship (#8386); Fabrizio A. Pennacchio by AIRC fellowship #23966. Work in Ruocco’s group was partially funded by grants from ERC-2019-Synergy Grant (ASTRA, n. 855923); EIC-2022-PathfinderOpen (ivBM-4PAP, n. 101098989); Project “National Center for Gene Therapy and Drugs based on RNA Technology” (CN00000041) financed by NextGeneration EU PNRR MUR—M4C2—Action 1.4—Call “Potenziamento strutture di ricerca e creazione di “campioni nazionali di R&S” (CUP J33C22001130001). Work in the Zippo group was supported by Telethon Foundation (GGP20010); the European Innovation Council through its Horizon Europe Pathfinder Program under Grant Agreement No. 101098989; the Italian Association for Cancer Research (AIRC), IG# 22911; by the European Union under NextGenerationEU (PRIN 2022 #2022H79275).

## Author Contributions

S.D. and A.Z. conceived the study, designed the experiments, and interpreted the data. S.D., D.M., G.V., S.L., and C.B. performed the cellular and molecular biology studies, participating in data analyses. L.S. performed the in vivo studies and together with S.D. performed the tissue sectioning and staining. C.T., E.P., and G.R. performed the Brillouin microscopy and computational data analysis. A.P., F.P., and P.M provided fundamental materials and technologies for the study. E.S. and D.G. provided the biological samples and isolated primary fibroblasts. L.P., T.T., R.M:, and R.P. conceived and performed the molecular dynamics simulations. A.Z. supervised the work and wrote the manuscript.

## Methods

### Cell lines and cell culture conditions

The cell lines used in this study were hTERT-immortalized human adipose-derived MSCs (a gift from P. Tatrai) carrying frameshift mutation in the exon 39 of KMT2D, i.e. MLL4^Q4092X^ and MLL4^P4093X^ (MLL4^LoF_1^ and MLL4^LoF_2^ respectively)^10^, human primary fibroblasts derived from either healthy or Kabuki syndrome individuals (kindly provided by D. Genevieve), and NIH3T3 cells (American Type Culture Collection). MSCs were cultured in 1:1 DMEM/F-12 medium (Gibco; 11320-074) supplemented with 10% fetal bovine serum (Euroclone; ECS0180L) and 100 U/mL Penicillin/Streptomycin (Gibco; 15140122). Primary fibroblasts and NIH3T3 were maintained in DMEM medium (Gibco; 11965092) supplemented with 10% fetal bovine serum and 100 U/mL Penicillin/Streptomycin. Cells were maintained at 37 °C under 5% CO_2_. When needed, cells were treated with the STINGi (H-151 HY-112693) 0.2 µM for 24 hours or the STING agonist (diABZI HY-112921B) 0.1 µM for 16 hours. To assess the effect of different substrate stiffness on MSCs, we used the CytoSoft® Imaging 24-well Plates (Advanced Biomatrix; 5188-1EA) of 0.5 and 32kPa stiffness. Plates were washed three times with PBS and then coated with 10 µg/ml fibronectin (Santa Cruz; sc-29011) for 1h at RT. 20.000 cells/cm^2^ were plated and grown for 48h prior to analysis. For the optogenetics experiments, MSCs expressing piggyBAC-MLL4-PrLD-HaloTag-Cry2 were stained with 200nM of 646 Janelia Fluor® HaloTag® (Promega; GA1120) diluted in complete culture medium for 20 minutes at 37°C. The substrate was then quickly washed once in PBS before replacing the medium and performing time-lapse imaging.

To assess the level of rupture and cell death MSCs were plated in 35mm Petri dish containing a set of molded microchannels, with variable dimensions and restrictions (4DCell; MC019). Microchannel dishes were washed three times with PBS for 5 minutes. The last washing was replaced in each chamber by 10µl of 10µg/µl fibronectin and incubated for 1h at RT. After the coating, the device was washed three times with PBS, and the MSC culture medium containing 2% FBS was added to cover each chamber and left for 15 minutes at 37°C. Afterwards cells (1500.000 cells/10µl in 2% FBS-containing medium) were loaded in each access port and kept for 30 minutes at 37°C before adding 2ml of 2% FBS culture medium to the dish. Cells were grown for 48h. Six hours before live imaging, a gradient of FBS was created by adding a droplet of 10% FBS-containing medium in the access port closest to where the cells have been plated and with whom they communicate. Finally, 10% FBS-containing medium was added to fill each chamber and time-lapse video microscopy was performed.

To confine cells at 3 and 6 μm, the CSOW 620 – static confiner (4DCell) was used. Pistons and confinement slides were equilibrated for 1h at 37°C in the culture medium before performing live-cell imaging. To ensure cell adherence, the glass bottom 6-well plate was coated with 10 µg/µl fibronectin for 1h at RT, before plating 20.000 MSCs in the central circular part of the well. When needed, cells were stained with Hoechst and SiR-actin (Spirochrome; SC001). Confinement was performed 24h after plating.

To study the NE tensional state, MSCs expressing pCDH-MiniNesprin1-GFP-cpstFRET were seeded at a density of 20.000 cells/cm^2^ in an IBIDI plate (82426) and stained with SirDNA (Spirochrome; SC007). Imaging was performed after 48h. pCDH-MiniNesprin1-GFP-cpstFRET is an orientation-based FRET biosensor (circularly permutated stretch sensitive -cpst) that uses the N-terminus and C-terminus of Nesprin-1 protein as backbone. It was designed as described in Poli A.et al., ^47^ to be modulated by the angles between the donor (cpst Cerulean) and acceptor (cpst Venus).

### Derivation of stable cell lines

MSCs expressing pTRIP-SFFV-EGFP-NLS (Addgene #86677), pCDH-MiniNesprin1-GFP-cpstFRET (gifted by Maiuri laboratory), pCDH–EF1–MCS–IRES–PURO–H3.3/K27M (gifted by Allis laboratory.), TETO-FUW-LMNA-HALO (gifted by Zaret laboratory), pTRIP-GFP-IRF3 (Addgene #127663), pTRIP-SFFV-cGAS-EGFP (Addgene #127661), pTRIP-CMV-mCherry-53BP1 (Addgene #127658), pTRIP-BAF-mCherry were obtained through transduction of MSCs with the corresponding lentiviral vectors. MSCs stably expressing piggyBAC-MLL4-PrLD-HaloTag-Cry2 were obtained by co-nucleofecting 50.000 cells with piggyBAC-MLL4-PrLD-HALOTag-Cry2 (1µg) and pCMV-Pbase (0.2µg) (kindly provided by L. Tiberi) using P1 Primary Cell 4D-NucleofectorTM X Kit L (Lonza; V4XP-1024) Amaxa Nucleofector (program FF104, Lonza) following manufacturer’s instructions.

### DNA constructs

For the optogenetic experiments, the construct piggyBAC-MLL4-PrLD-HaloTag-Cry2 was cloned as follows: the HaloTag sequence was PCR amplified and then cloned in the piggyBAC mCherry-MLL4-PrLD-Cry2^10^, substituting the mCherry construct.

To overexpress BAF, the lentiviral vector pTRIP-mCherry-BANF1 was generated as follows: the BANF1 sequence was PCR amplified from pHM-Hyg-1xFlag-BAF-WT (a kind gift from Matthew S. Wiebe). The fragment was then cloned in the lentiviral vector pTRIP-mCherry-53BP1 substituting the 53BP1 construct. Oligonucleotides used for PCR amplification of cloning fragments are listed in Supplementary Table 1.

### Immunofluorescence

Cells were fixed with 4% PFA for 10 min at room temperature (RT) and washed three times with phosphate-buffered saline (PBS). Immunostaining was performed directly on the CytoSoft® Imaging 24-well Plates as follows: cells were permeabilized and blocked with PBS/ 1% bovine serum albumin (Millipore; 126579)/ 5% goat serum (Fisher Scientific; 11475055)/ 0.5% Triton X-100 (blocking solution) for 1 h at room temperature. Cells were incubated overnight at 4 °C with primary antibodies diluted in the blocking solution, washed in PBS three times, and incubated for 1h at RT with secondary antibodies (Alexa Fluor conjugated produced in goat by Thermo Fisher) diluted in the blocking solution, comtaining DAPI (Sigma–Aldrich; D9542) for nuclear staining. After three washings with PBS, the mounting media ProLong Gold (Invitrogen; P36934) was added to each well and then covered with coverslips to preserve the staining. The primary antibody and corresponding dilutions are listed in Supplementary Table 2.

Confocal images were acquired using a Leica TCS SP8 microscope with a HCX Plan Apo ×63/1.40 objective. When needed (nuclear volume and flatness quantification), z stacks were acquired with sections of 0.5 μm. Image acquisition settings were kept constant for downstream image analysis. SIM images were acquired using a Nikon Eclipse Ti2E N-SIM microscope in 2D using a CFI Apo SR HP TIRF 100XC/1.49 objective, with ROI of 512×512 and 0.2 μm Z stacks.

### Live-cell Imaging

Time-lapse video microscopy was performed at 37 °C and 5% CO_2_ using the Nikon Eclipse Ti2 with an ×20/0.75, ×60/1.4 or ×100/1.45 Plan Apo λ objective (Nikon) integrated with a Lumencor SpectraX LED light source system and a EMCCD sensor camera (Andor iXon Ultra 888) for the detection. When imaging cells in microchannels, each frame was acquired every 5-10 minutes for at least 12h. When imaging cells under confinement, images were taken every 10 minutes for 2-10 hours, starting the acquisition 20 minutes after applying the confiner. To avoid fluorophore bleaching and cell phototoxicity during time-lapse, imaging LED intensity and exposure time were kept between 4-25% and 5-100 ms, respectively. For the optogenetics experiments time-lapse video was carried out continuously for the indicated timings under controlled temperature and CO_2_. Images of fluorescent cells were acquired before and after the stimulus (blue-light activation with LED 470 100% intensity for 3 seconds). and then every 30 seconds. MSCs expressing pCDH-MiniNesprin1-GFP-cpstFRET were acquired using Leica TCS SP8 confocal microscope with a HCX Plan Apo ×63/1.40 objective. Donor (cpCerulean) was excited at 458 nm, and emission peaks of cpCerulean and cpVenus were captured, respectively, in a window of 470-490nm and 520-540nm.

### Confocal Brillouin microscopy, data acquisition and analysis

The Confocal Brillouin Microscope consists of an inverted Olympus IX-73 coupled to a double virtually imaged phased array (VIPA)-based spectrometer through optical fibers. The laser source (λ=532 nm, VERDI) is focused on the sample plane via a Olympus UPlanSApo 60x/ 1.4 NA objective to have high resolution Brillouin shift maps of biological samples: an higher Brillouin shift is correlated with higher local stiffness, with a spatial resolution comparable to confocal ones (∼400 nm on xy plane, ∼800 nm on z). The 3D mechanical properties of the sample can be recovered in confocal laser scanning mode thanks to a pair of galvanometric mirrors (THORLABS) and a piezo stage (MadCity Labs). Standard brightfield and fluorescence units are located adjacent to the Brillouin laser-scanning module, allowing for the simultaneous comparison of morphological, fluorescent and mechanical data of any biological sample. The Microscope setup is controlled in MATLAB via a custom-built GUI.

For the analysis of the nuclear mean Brillouin shift in cells and tissues stained with DAPI, a custom-made MATLAB program created a binary mask from fluorescence data and calculated the mean Brillouin shift across the masked region of each nucleus. For the analysis of the ECM mean Brillouin shift in tissues, another custom-made MATLAB code automatically segmented different regions of the maps, selected all the pixels having Brillouin shifts values above 7.80 GHz (excluding the pixels corresponding to fluorescence masked regions), connected different parts of the images with the same shift and calculated the mean value across these regions.

### Image analysis

Images acquired by confocal were analyzed using ImageJ (FIJI) software.

For 2D analysis, the DAPI signal was used to define the ROI (region of interest) of the nucleus. To measure the nuclear mean intensity and area, LIF files were firstly converted to TIFF composite images, then the minimum value of the threshold and the size range parameters were determined for each independent experiment to identify the nuclei on the binary image. For the measure of nuclear volume and flatness, a 3D analysis was performed by using the 3D plugin suite (ImageJ plugin) and segmenting the nuclei on the DAPI signal. For measuring cluster size, mean intensity, and area in 2D the analysis was performed in ImageJ as follows: first, background subtraction was applied (rolling ball correction), then unsharp masking and median filters were carried out. The clusters were finally identified with the Shanbhag dark automatic threshold.

For calculating the Excess of Perimeter (EOP), the ratio of internal and perimetral length of the NE (based on Lamin B staining) over the perimeter of the convex envelope was measured using in-house ImageJ macro as previously published^47^.

The percentage of cGAS-positive cells was measured by counting the number of cells showing cGAS accumulation at the nuclear periphery over the total number of cells acquired. The percentage of BAF-positive cells was measured by counting the number of cells showing BAF clustering (pTRIP-BAF-mCherry MSCs) over the number of cGAS-positive cells. The percentage of BAX-activated cells was measured by counting the number of cells showing BAX signal matching the MitoTracker signal over the number of the cGAS-positive cells (pTRIP-SFFV-cGAS-EGFP MSCs). The percentage of cells with nuclear IRF3 (pTRIP-GFP-IRF3 MSCs) was measured by counting the number of cells showing accumulation of IRF3 in the nucleus over the number of the cGAS-positive cells. For the microchannel experiments, image analysis and single-cell tracking were carried out using the NIS software. Nuclear Envelope Rupture was quantified on the basis of the NLS-EGFP signal as described in a previous work^16^. For measuring the NLS-EGFP Intensity in the cytoplasm and nucleoplasm, a small ROI was put in front of the nucleus at each time frame in which the nucleus enters and passes through the restriction (ROI corresponding to the cytoplasm). The average intensity of the ROI of the cytoplasm was divided by the average intensity of the nuclear NLS-GFP signal before the nucleus entered the constriction (Cytoplasm/Nucleoplasm ratio) or vice-versa (Nucleoplasm/Cytoplasm ratio). The percentage of rupture was calculated as the ratio between the number of cells undergoing nuclear envelope rupture and the number of cells passing through the constriction. The percentage of cell death was measured by calculating the ratio between the number of cells dying while passing through the restriction and the total number of cells passing through the constriction. The percentage of cells with cGAS, BAF, 53BP1, and MLL4-PrLD foci was calculated based on the number of cells showing rounded clusters of the corresponding marker while passing through restrictions over the total number of cells that passed.

For the optogenetics experiments the images of fluorescent cells were analyzed to obtain the number, area, and mean intensity of clusters as follows: maximum intensity projection in z was performed (z-step size of 0.5 μm), then the ROI was drawn based on the HaloTag 646 fluorescent intensity signal on the pre-stimulus condition. After background subtraction, in which the rolling ball correction was kept constant for every experiment, the following threshold was used to identify clusters: “(mean intensity of the nucleus on the pre-stimulus) + (4X standard deviation)”. This allowed us to consider variations in the expression of the protein before the stimulus at the single-cell level. The function “Find Maxima” with a prominence of 20 was used to further define the clusters.

The inverted FRET index was calculated as the ratio between cpCerulean and cpVenus mean intensity. Data were analyzed using the SirDNA channel to define nuclear ROI. For the representative images, the PixFRET ImageJ plug-in was used to retrieve FRET efficiency maps and the out-of-the-nucleus background was subtracted.

### Animal studies

Mice were housed in a certified animal facility in accordance with European Guidelines. The experiments were approved by the Italian Ministry of Health as conforming to the relevant regulatory standards. *Kmt2d^fl/fl^* mice were a kind gift from I. Pastushenko and were previously generated and described to target the SET domain of Mll4^42^. CMV-Cre mice (JAX #006054) were purchased from Charles River Laboratories. Control strain (Kmt2d*^fl/+^*) was generated crossing *Kmt2d^fl/fl^* with C57BL/6J mice. KS mouse model (Kmt2d*^cHet^*) was obtained crossing *Kmt2d^fl/fl^* with CMV-Cre mice to obtain Mll4 haploinsufficiency. Mouse body growth and length were monitored at selected time points from postnatal day 14 (P14) to P30. To obtain the measures of mouse body length, mice were anesthetized with 5% Isoflurane and recorded with a camera. A single frame was isolated in post-processing and body length from nose to tail was measured using ImageJ (FIJI) software. Animals were sacrificed at P30 for subsequent histological analysis.

### Histological examination of mouse tissues

Mice were euthanized using CO_2_ and cervical dislocation. Tibiae and femurs were collected and fixed with 4% PFA for 48h. Tissues were cryopreserved in 30% sucrose overnight, embedded in Frozen Section Compound (Leica; 3801480) and sectioned at 30 µm using Thermo Scientific HM525 NX cryostat. For immunofluorescence, frozen sections were initially stained with Phalloidin-TRITC to visualize f-actin, post-fixed with 4% PFA and then incubated with 10 mM sodium citrate, pH 6.0 for 20 minutes at 95°C. Incubation with blocking solution, primary and secondary antibodies were performed as previously described for the *in vitro* analysis. Sections were mounted using aqueous mounting medium and acquired using a Leica TCS SP8 confocal microscope with HC Plan Apo ×20/0.75 or HCX Plan Apo ×63/1.40 objective. For H&E staining and histological analysis samples were acquired with the Zeiss Imager M2 microscope with ZEISS Axiocam 705 color camera and EC Plan-Neofluar ×20/0.5 objective. For femurs, longitudinal growth plate sections were used, and proliferative zone, hypertrophic zone, and total growth plate heights were measured using ImageJ (FIJI) software in at least 6 sites per section (1 section per mouse).

### Molecular dynamics simulation

Atomistic MD simulations of the extended polyglutamine (polyQ) tract belonging to the intrinsically disordered, Prion-like domain of MLL4 (MLL4-PrLD)^10^ were carried out with the Gromacs 2020.3 software, in combination with the a99SBdisp force field/TIP4P explicit water model^63^. Putative coordinates of MLL4-PrLD were obtained via the [IntFold web server](https://www.reading.ac.uk/bioinf/IntFOLD/), whereby we extracted the structure of the polyQ stretch (i.e., residues 338 to 414). The latter was solvated within a rhombic dodecahedral box and neutralized by an excess concentration of monovalent salt [KCl] = 0.15 M. Periodic boundary conditions in the three dimensions were associated with the Particle Mesh Ewald calculation of the full-system electrostatics, and a 12 Å cutoff was established to both Van der Waals and short-range electrostatic forces. In addition, LINCS constraints were applied to all hydrogen atoms, thereby allowing for an integration timestep of 2 femtoseconds.

Prior to raw data collection, a minimization/thermalization protocol has been followed that involves i) 5000 minimization steps by a steepest descent algorithm, ii) a 0.5-nanosecond thermalization of the solvent bath in the NVT ensemble at T = 310 K, and iii) a 5-nanosecond further equilibration in the NPT ensemble at P = 1 atm - the last two stages restraining the coordinates of the solute. Fixed T, P were enforced by a stochastic velocity-rescale thermostat (t = 0.1 ps) and an isotropic Parrinello-Rahman barostat (t = 2.0 ps) respectively. Thus, subsequent 1-microsecond MD simulations of the polyQ tract at increasing target pressures - i.e. 1, 1.25, 1.5 and 1.75 atm - have followed, each trajectory starting from the last frame of the preceding stage on the pressure “ramp”. Yet, to ensure the stability and convergence of the system, each stage has been preceded by a 5-nanosecond equilibration of the solvent bath in the NPT ensemble - as done similarly in the aforementioned protocol. Lastly, raw data analyses and graphical rendering of the MD trajectories were performed with the [VMD software](https://www.ks.uiuc.edu/Research/vmd/), whereas scientific Python libraries were employed for the elaboration and plotting of data.

### Statistical Analysis

No statistical method was used to predetermine sample size. All experiments were performed on at least three independent biological samples and the relative measurements were taken from distinct replicates. For quantification of imaging data, all of the images were acquired by random sampling through acquisition of at least 10 non-overlapping fields of view/condition. The acquisition and the relative quantifications were reproduced using biologically independent samples, retrieving similar results. The exact statistical parameters used for each experiment are reported in figure captions. The statistical tests performed are two-tailed unpaired Student’s t-test and two-way ANOVA for multiple comparisons. P-values are indicated in the figures and the corresponding figure captions as: *= p<0.05, **= p<0.01, ***= p < 0.001, ****= p < 0.00001, ns = not significant (p<0.05). Data collection and analyses of all studies involving animals were conducted randomly without blinding.

