## Supplementary Figures for "Chromatin condensates tune nuclear mechano-sensing in Kabuki Syndrome by constraining cGAS activation"

### Extended Data

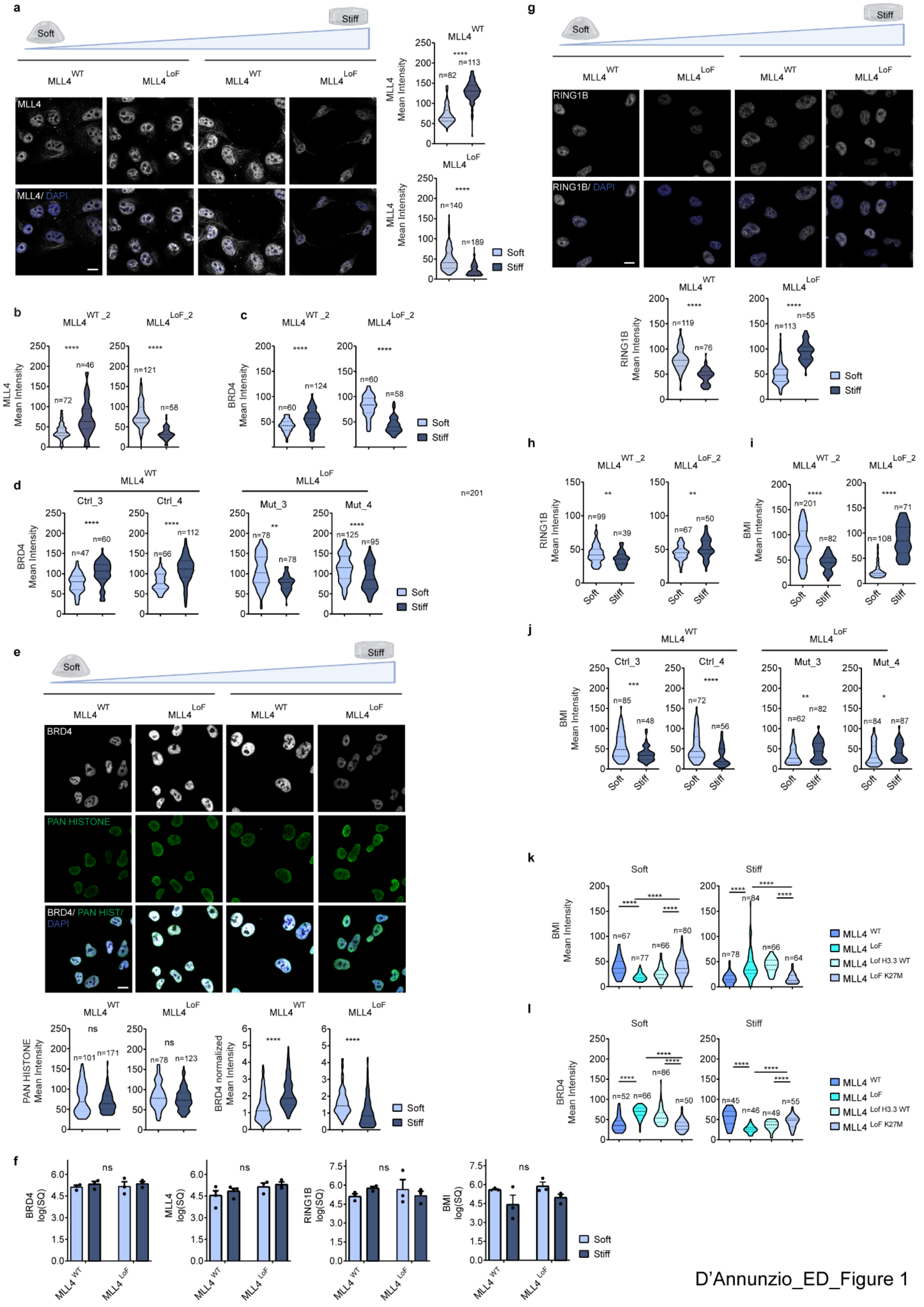


D’Annunzio_ED_Figure 1

**Extended Data Fig. 1**

**(a)** Representative confocal images and relative quantification of nuclear MLL4 mean intensity in MLL4$^{WT}$ and MLL4$^{LoF}$ MSCs on soft and stiff matrix. Scale bars, 10 µm. **(b)** Quantifications of nuclear MLL4 mean intensity in MLL4$^{WT_2}$ and MLL4$^{LoF_2}$ MSCs independent clones on soft and stiff matrix. **(c)** Quantification of nuclear BRD4 mean intensity in MLL4$^{WT_2}$ and MLL4$^{LoF_2}$ MSCs independent clones on soft and stiff matrix. **(d)** Quantification of nuclear BRD4 mean intensity in primary fibroblasts from healthy donor (Ctrl) or Kabuki patients (Mut) on soft and stiff matrix. **(e)** Representative confocal images and relative quantifications of nuclear BRD4 and PAN HISTONE mean intensity in MLL4$^{WT}$ and MLL4$^{LoF}$ MSCs on soft and stiff matrix. Scale bars, 10 µm. **(f)** Real-time quantitative reverse transcription PCR of the indicated genes in MLL4$^{WT}$ and MLL4$^{LoF}$ MSCs on soft and stiff matrix ($n$ = 3 biologically independent samples). **(g)** Representative confocal images and relative quantifications of immunostaining for RING1B in MLL4$^{WT}$ and MLL4$^{LoF}$ MSCs on soft and stiff matrix. Scale bars, 10 µm. **(h)** Quantifications of nuclear RING1B mean intensity in MLL4$^{WT_2}$ and MLL4$^{LoF_2}$ MSCs independent clones on soft and stiff matrix. **(i)** Quantifications of immunostaining for BMI in MLL4$^{WT_2}$ and MLL4$^{LoF_2}$ MSCs independent clones on soft and stiff matrix. **(j)** Quantifications of nuclear BMI mean intensity in primary fibroblasts from healthy donor (Ctrl) or Kabuki patients (Mut) on soft and stiff matrix. Quantifications of immunostaining for BMI **(k)** and BRD4 **(l)** in MLL4$^{WT}$ and MLL4$^{LoF}$ MSCs, as well as MLL4$^{LoF}$ MSCs expressing either H3.3WT or H3.3K27M on soft and stiff matrix. Violin plots in **(a)**- **(e)** and **(g)**- **(l)** indicate median values (middle lines), and first and third quartiles (dashed lines). Bar plots in **(f)** show mean + S.E.M. The number of cells ("n") analyzed is reported in each panel. Statistical significance was determined by a two-tailed unpaired student's t-test.

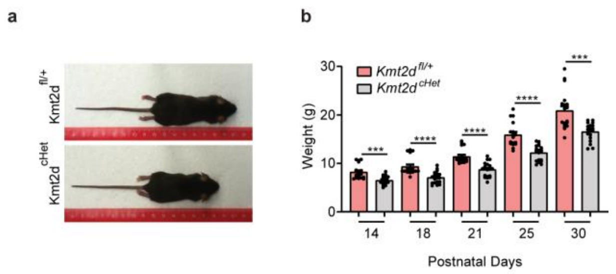


D’Annunzio_ED_Figure 2

**Extended Data Fig. 2**

**(a)** Representative images of Kmt2d$^{fl/+}$ (WT) and Kmt2d$^{cHet}$ (KS). **(b)** Quantification of mice weight (g) in Kmt2d$^{fl/+}$ (WT) and Kmt2d$^{cHet}$ (KS) at the indicated postnatal days (*n* = 18 mice derived from three different litters). Bar plot in **(b)** shows mean + S.E.M.. Statistical significance was determined by a two-tailed unpaired student's t-test.

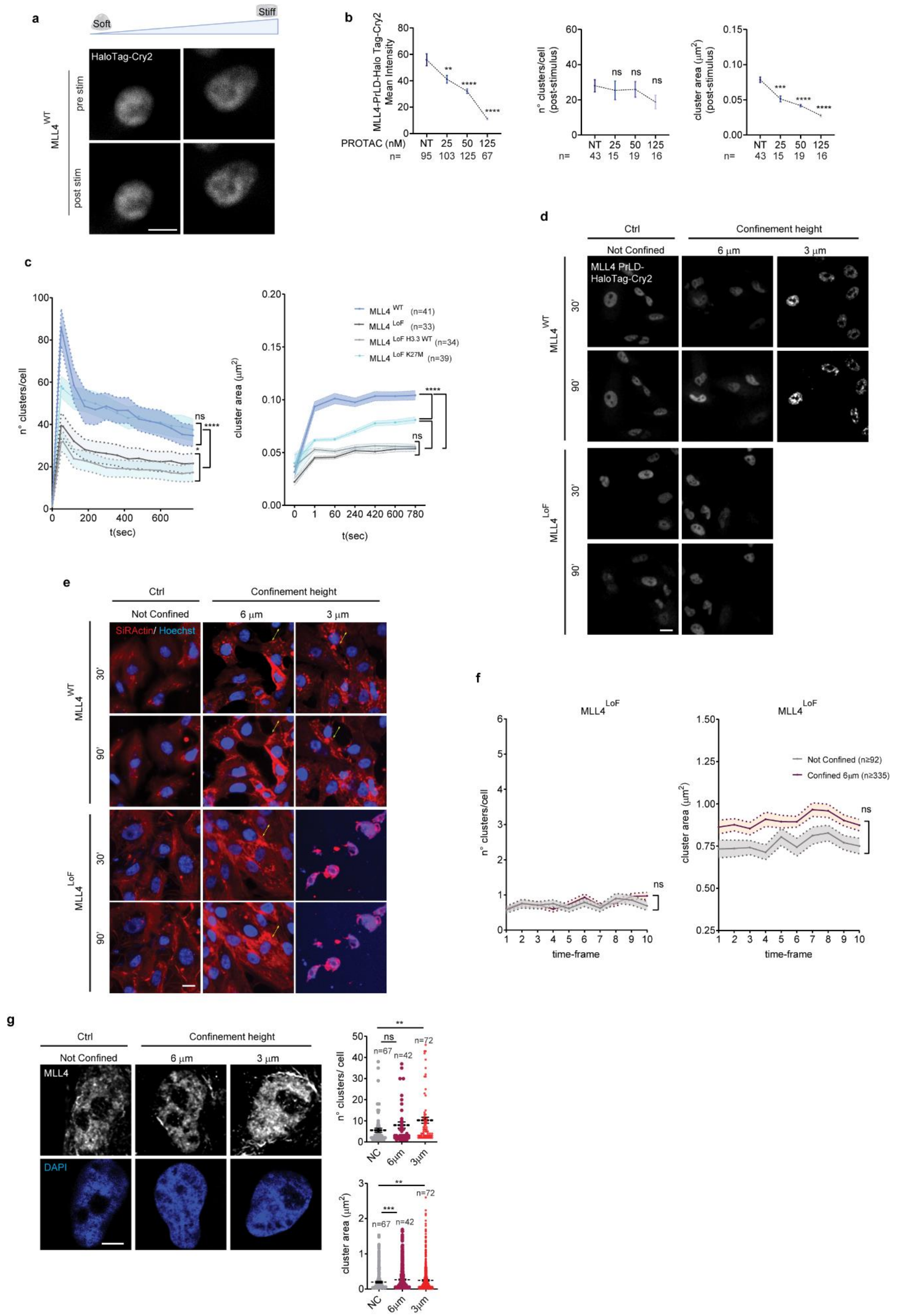


D'Annunzio_ED_Figure 3

35 **Extended Data Fig. 3**

**(a)** Representative images of MSCs on soft and stiff matrix over-expressing HaloTag-Cry2 before and after the stimulus with blue light. Scale bars, 10 µm. **(b)** Quantification of the number and area of light-induced droplets of MLL4-PrLD-HaloTag-Cry2 in MLL4$^{WT}$ MSCs after the blue light stimulus treated with PROTAC3 at the concentration indicated (NT=Non Treated). **(c)** Quantification of the number and area of light-induced droplets of MLL4-PrLD-HaloTag-Cry2 at different time points in MLL4$^{WT}$ and MLL4$^{LoF}$ MSCs, as well as MLL4$^{LoF}$ MSCs expressing either H3.3WT or H3.3K27M on soft and stiff matrix. The time point t=0 represents the pre-stimulus. **(d)** Representative image of MLL4$^{WT}$ and MLL4$^{LoF}$ MSCs over-expressing MLL4-PrLD-HaloTag-Cry2 not confined and confined at 6 and 3µm at the indicated timing. Scale bars, 10 µm **(e)** Representative images showing MLL4$^{WT}$ and MLL4$^{LoF}$ MSCs not confined, and under confinement of 6 and 3µm at the indicated time points. Cell nuclei are stained with Hoechst (blue), whether the cytoskeleton is marked in red (SirActin). Yellow arrows indicate actomyosin recruitment to the cell-cortex in response to confinement. Scale bars, 10 µm. **(f)** Quantification of the number and area of confinement-induced droplets of MLL4-PrLD-HaloTag-Cry2 at different time points in MLL4$^{LoF}$ MSCs . **(g)** Representative confocal images and relative quantifications of immunostaining for MLL4 in MLL4$^{WT}$ MSCs not confined, and confined at 6 and 3µm for 45 minutes. Scale bars, 10 µm. Scatter XY plots in **(b)**, **(c)** and **(f)** and scatter dot plots in **(g)**- **(h)** show mean + S.E.M. The number of cells ("n") analyzed is reported in each panel. Statistical significance was determined by two-way ANOVA test for panels **(c)** and **(f)**, and by a two-tailed unpaired student's t-test for panel **(g).**

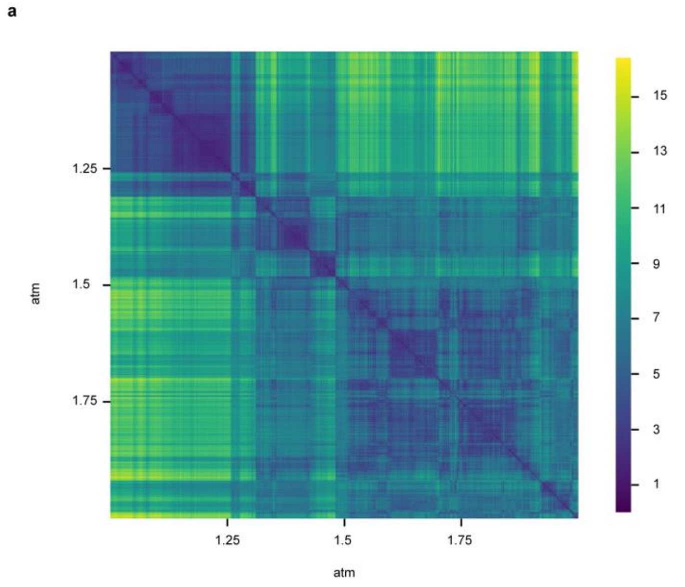


D'Annunzio_Figure_ED4

**Extended Data Fig. 4**

**(a)** Two-dimensional root mean square displacement (RMSD) profile of the polyQ tract of MLL4-PrLD, based upon the coordinates of the two most stable helices along the MD trajectory (i.e., conserved in over 75% of the trajectory frames and involving residues 3928-3935 and residues 3948-3954). A low value of the RMSD is associated with a high degree of structural/conformational similarity between frames: Here, two broad squares (blue shades) might be identified that corroborate the existence of a twofold conformational basin of the polyQ tract, the higher pressure regimes stabilizing a more expanded configuration. Analysis performed within the MD Analysis framework (PMID: 15973002, 21500218).

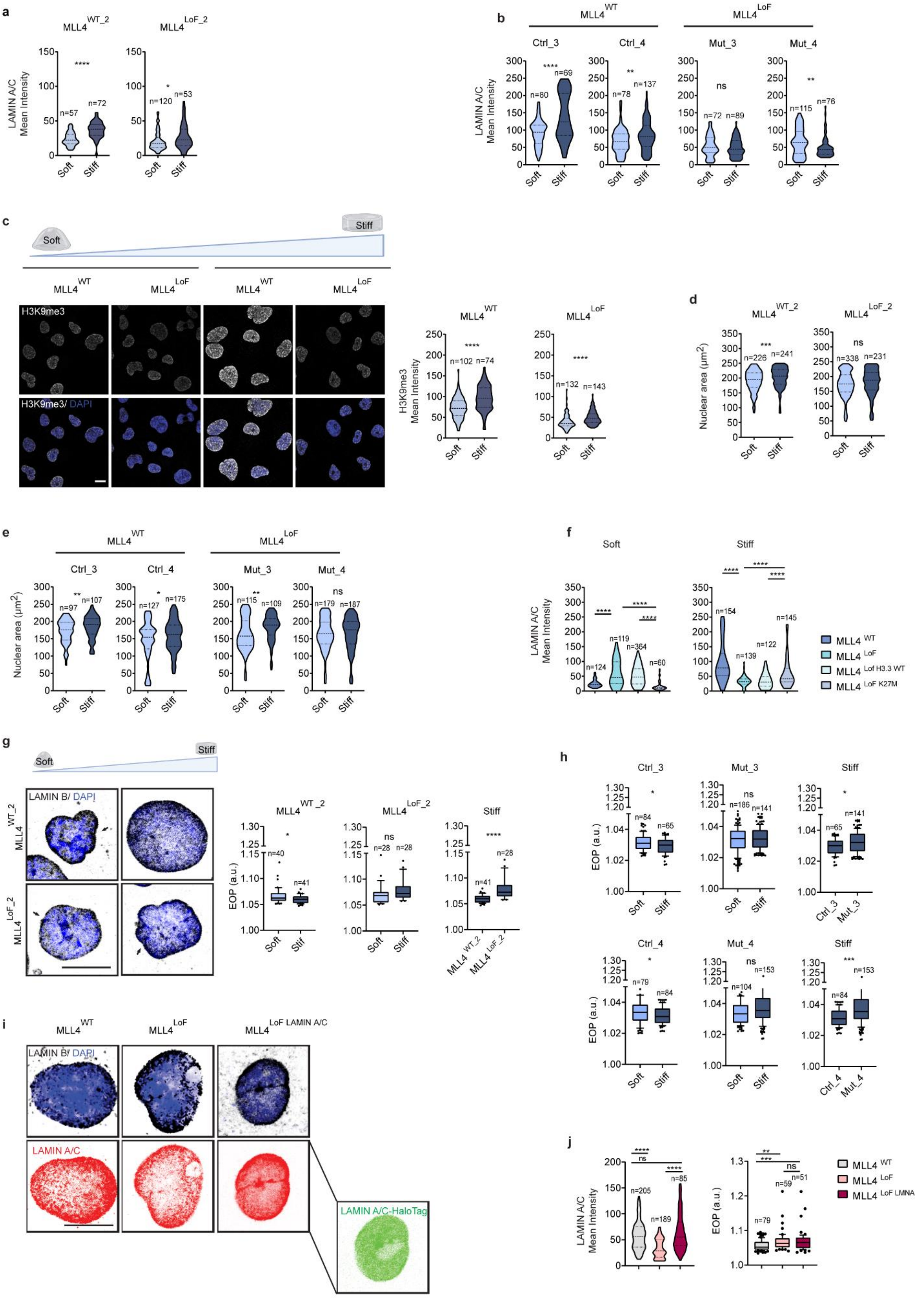




D'Annunzio_ED_Figure 5

75 **Extended Data Fig. 5**

**(a)** Quantifications of immunostaining for LAMIN A/C in MLL4$^{WT_2}$ and MLL4$^{LoF_2}$ MSCs independent clones on soft and stiff matrix. **(b)** Quantifications of nuclear LAMIN A/C mean intensity in primary fibroblasts from healthy donor (Ctrl) or Kabuki patients (Mut) on soft and stiff matrix. **(c)** Representative confocal images and relative quantifications of immunostaining for H3K9me3 in MLL4$^{WT}$ and MLL4$^{LoF}$ MSCs on soft and stiff matrix. Scale bars, 10 μm. Quantifications of nuclear area in MLL4$^{WT}$ and MLL4$^{LoF}$ MSCs independent clones **(d)** and in primary fibroblasts **(e)** on soft and stiff matrix. **(f)** Quantifications of immunostaining for LAMIN A/C in MLL4$^{WT}$ and MLL4$^{LoF}$ MSCs as well as MLL4$^{LoF}$ MSCs expressing either H3.3WT or H3.3K27M on soft and stiff matrix**.** Representative confocal images and relative quantification of immunostaining for LAMIN B in MLL4$^{WT}$ and MLL4$^{LoF}$ MSCs independent clones **(g)** and primary fibroblasts **(h)** on soft and stiff matrix. Black arrows indicate the presence of nuclear envelope invaginations, quantified as EOP (Excess Of Perimeter) ($n \geq 2$ biological replicates). Scale bars, 10 μm.

**(i)** Representative confocal images of immunostaining for LAMIN B and LAMIN A/C in MLL4$^{WT}$ and MLL4$^{LoF}$ MSCs as well as MLL4$^{LoF}$ MSCs expressing Halo-tag LAMIN A/C. **(j)** Box-plots showing quantification of EOP in MLL4$^{WT}$, MLL4$^{LoF}$ and MLL4$^{LoF}$ expressing Halo-tag LAMIN A/C MSCs. Scale bars, 10 μm.

Violin plots in **(a)**- **(c)** and **(j)** indicate median values (middle lines), and first and third quartiles (dashed lines). Box plots in **(g)**- **(j)** indicate the median (middle line), the first and third quartiles (box), and the 10th and 90th percentile (error bars) of the EOP. The number of cells ("n") analyzed is reported in each panel. Statistical significance was determined by a two-tailed unpaired student's t-test.

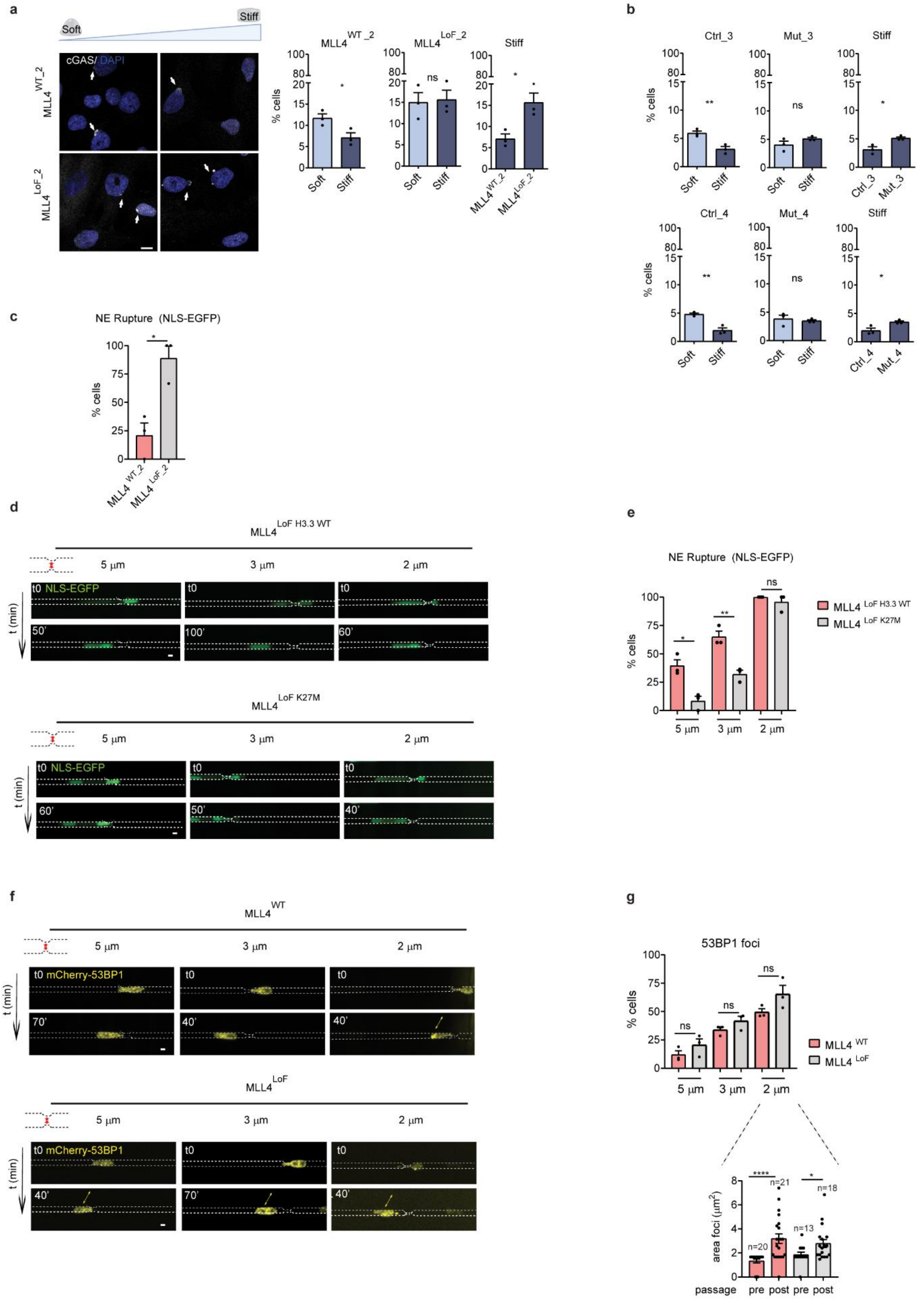




D'Annunzio_ED_Figure 6

**Extended Data Fig. 6**

**(a)** Representative confocal images of immunostaining for cGAS in MLL4$^{WT}$ and MLL4$^{LoF}$ MSCs independent clones on soft and stiff matrix. Scale bars, 10 µm. On the right, quantification of the percentage of cells showing cGAS accumulation at the nuclear periphery, indicating NE rupture (white arrows) (n=3 biologically independent samples). **(b)** Quantification of the percentage of cells showing cGAS accumulation at the nuclear periphery in primary fibroblasts from healthy donor (Ctrl) or Kabuki patients (Mut) on soft and stiff matrix , indicating NE rupture (white arrows) (n=3 biologically independent samples). **(c)** Quantification of the percentage of cells undergoing NE rupture in NLS EGFP-MSCs (independent clones) (n=3 biologically independent samples)**. (d)** Representative images of MLL4$^{LoF}$ H3.3WT and H3.3 K27M - NLS-EGFP in the process of passing through restrictions of different widths at the indicated time points. Scale bars, 10 µm. On the right, Bar plots showing the percentage of cells undergoing NE rupture (n=3 biologically independent samples) **(e). (f)** Representative images of MLL4$^{WT}$ and MLL4$^{LoF}$ MSCs expressing mCherry 53BP1 passing through constrictions of different widths (5, 3 and 2 µm). Scale bars, 10 µm. **(g)** Quantification of the percentage of cells harboring 53BP1 foci (n=3 biologically independent samples)**.** Below, bar plots showing the area of the mCherry 53BP1 cluster before and after passage through restrictions of 2 µm (n=3 biologically independent samples). The number of cells ("n") analyzed is reported in the figure. Bar plots show mean + S.E.M. Statistical significance was determined by a two-tailed unpaired student's t-test.

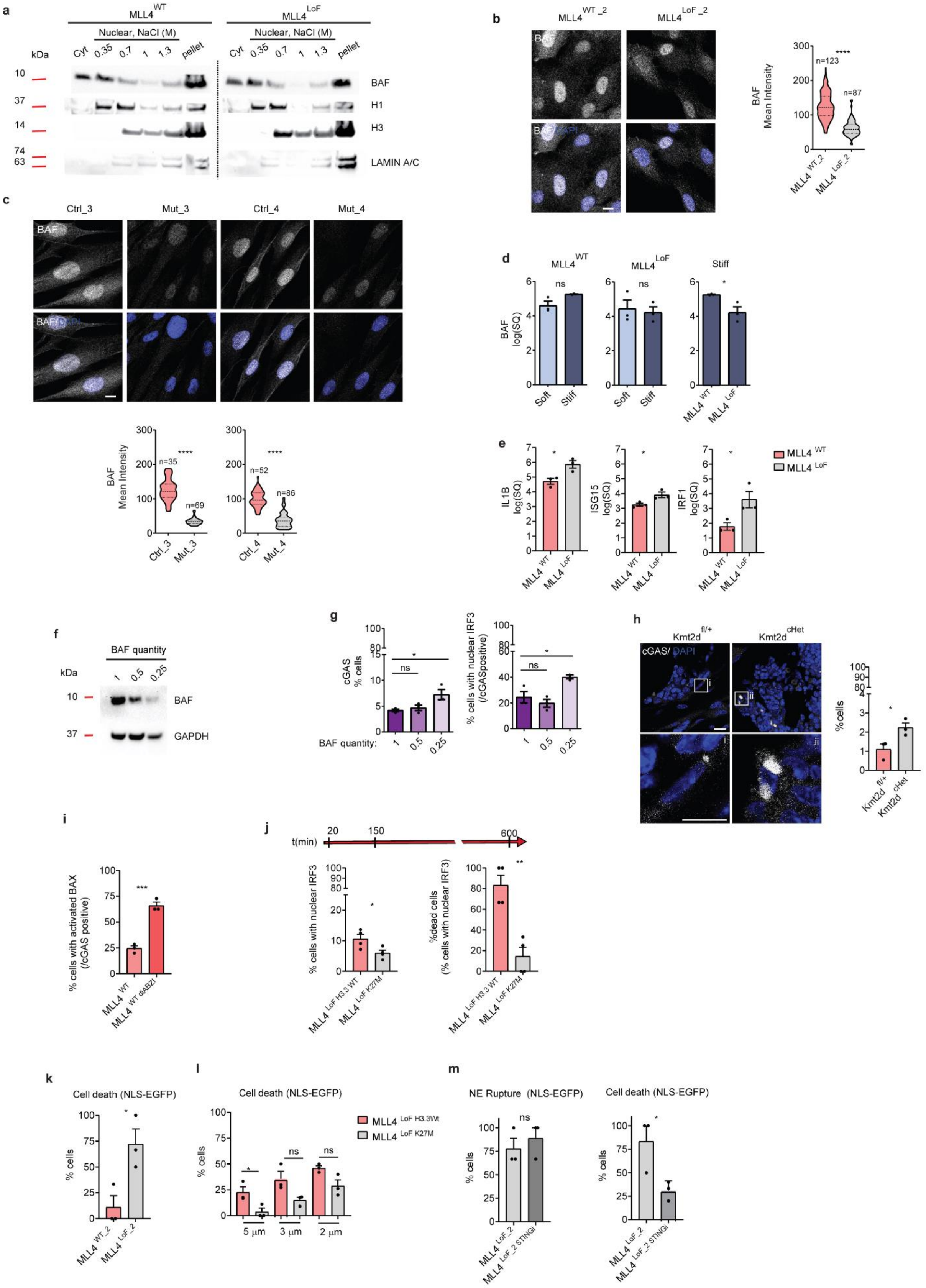




D'Annunzio_ED_Figure 7

123

**Extended Data Fig. 7**

**(a)** Western Blot analysis of BAF, Histone H1, Histone H3 and LAMIN A/C in MLL4$^{WT}$ and MLL4$^{LoF}$ MSCs extracts obtained using a gradient of NaCl. **(b)** Representative confocal images and e quantifications of nuclear BAF mean intensity in MLL4$^{WT_2}$ and MLL4$^{LoF_2}$ MSCs on the stiff matrix. The number of cells (“n”) analyzed is reported in the figure. Scale bar, 10 µm. **(c)** Representative confocal images and relative quantifications of immunostaining for BAF in primary fibroblasts from healthy donor (Ctrl) or Kabuki patients (Mut) on the stiff matrix. The number of cells (“n”) analyzed is reported in the figure. Scale bar, 10 µm. **(d)** Real-time quantitative reverse transcription PCR of BAF in MLL4$^{WT}$ and MLL4$^{LoF}$ MSCs on soft and stiff matrix ($n = 3$ biologically independent samples). **(e)** Real-time quantitative reverse transcription PCR of IL1B, ISG15 and IRF1 in MLL4$^{WT}$ and MLL4$^{LoF}$ MSCs on stiff matrix ($n = 3$ biologically independent samples). **(f)** Western Blot analysis of BAF in MLL4$^{WT}$ MSCs transduced at different MOI of sgRNA targeting BANF1 promoter to obtain a gradient of BAF quantity (1=SCRAMBLE, 0.5= MOI 1, 0.25= MOI 0.25). **(g)** Quantification of the percentage of cells showing cGAS accumulation at the nuclear periphery and harboring IRF3 nuclear localization (calculated over the cGAS positive cells) in MLL4$^{WT}$ expressing different level of BAF protein ($n = 3$ biologically independent samples). **(h)** Representative confocal images of immunostaining for cGAS in the primary spongiosa of Kmt2d$^{fl/+}$ and Kmt2d$^{cHet}$ mice. Scale bar, 10 µm. On the right, quantification of the percentage of cells showing cGAS accumulation at the nuclear periphery (n=3, data combined from three experimental groups). **(i)** Quantification of the percentage of cells with activated BAX measured over the cGAS positive cells in MLL4$^{WT}$ MSCs +/- treated with diABZI (n=3 biologically independent samples). **(j)** Quantification of the percentage of MLL4$^{LoF}$ H3.3WT and H3.3

K27M- IRF3-EGFP harboring IRF3 nuclear localization in the first 150' of time-lapse (6μm confinement). On those cells having nuclear IRF3 was calculated the percentage of cells that undergo cell death within 600' of time-lapse (n=4, biologically independent samples). **(k)** Quantification of the percentage of cells undergoing cell death in NLS EGFP- MSCs independent clones passing through restrictions of 3μm (n=3 biologically independent samples)**. (l)** Quantification of the percentage of cells undergoing cell death in $MLL4^{LoF}$ H3.3WT and H3.3 K27M - NLS-EGFP passing through restrictions of 3μm. (n=3 biologically independent samples). **(m)** Quantification of the percentage of cells undergoing NE rupture and cell death in NLS EGFP- $MLL4^{LoF}$ MSCs treated or not with the STINGi passing through restrictions of 3μm (n=3 biologically independent samples)**.** Bar plots in **(d)**, **(e)**, **(g)**, **(h)**- **(m)** show mean + S.E.M. Violin plots in **(b)** and **(c)** indicate median values (middle lines), and first and third quartiles (dashed lines). Statistical significance was determined by a two-tailed unpaired student's t-test.
